# Genome-Wide Associations with Resistance to Bipolaris Leaf Spot (*Bipolaris oryzae* (Breda de Haan) Shoemaker) in a Northern Switchgrass Population (*Panicum virgatum* L.)

**DOI:** 10.1101/424721

**Authors:** Kittikun Songsomboon, Ryan Crawford, Jamie Crawford, Julie Hansen, Jaime Cummings, Neil Mattson, Gary Bergstrom, Donald Viands

**Author notes:** Corresponding Author: K. Songsomboon, Section of Plant Breeding and Genetics, School of Integrative Plant Science, Cornell University, Ithaca, NY 14853, USA.

## Abstract

Switchgrass (*Panicum virgatum* L.), a northern native perennial grass, suffers from yield reduction from Bipolaris leaf spot caused by *Bipolaris oryzae* (Breda de Haan) Shoemaker. This study aimed for determining the resistant populations via multiple phenotyping approaches and identifying potential resistance genes to the disease from genome-wide association studies in the switchgrass northern association panel. The disease resistance was evaluated from both natural (field evaluations in NY and PA) and artificial inoculations (detached leaf and leaf disk assays). There are ten out of the 66 populations showed the most resistant based on a combination of detached leaf, leaf disk, and mean from two locations. The GWAS from five subgroups from the association panel to different disease evaluation combinations yielded 27 significant SNPs on 12 chromosomes: 1K, 2K, 2N, 3K, 3N, 4N, 5K, 5N, 6N, 7K, 7N, and 9N accumulatively explaining phenotypic variance of BLUPs of detached leaf percent lesion via image analysis 26.52% at most and BLUPs of leaf disk percent lesion via image analysis 3.28% at least. Within linkage disequilibrium of 20 kb, these SNP markers linked with the potential resistance genes including genes encoding for NBS-LRR, PPR, cell wall related proteins, homeostatic proteins, anti-apoptotic proteins, and ABC transporters.

## INTRODUCTION

Switchgrass (*Panicum virgatum* L.) is a perennial biomass crop native to North America. Biomass and other agronomic traits have been the focus of breeding aims (Godshalk et al., 1998; Hopkins et al., 1993; Talbert et al., 1983; Vogel et al., 2002). Although many diseases have been reported to cause deleterious effects on yield, research on disease resistance, especially breeding for the resistance, is scarce. Only 28 of the 1693 research articles were directly related to diseases in switchgrass based on AGRICultural OnLine Access (AGRICOLA) (2018). *Bipolaris oryzae* (Breda de Haan) Shoemaker (teleomorph: *Cochliobolus miyabeanus*) is one of the major fungi causing Bipolaris leaf spot (BLS) in switchgrass that can reduce biomass by 70% (Fajolu, 2012). To manage the disease, breeding is one of the most economical approaches (Casler, 2012).

The natural distribution of switchgrass is broad latitudes across North America east of the Rocky Mountains. By simple morphological differentiation, switchgrass can be separated into two groups of upland and lowland ecotypes. Upland ecotypes are well adapted to higher latitudes and have higher drought tolerance whereas lowland ecotypes provide higher yield and require more water (Stroup et al., 2003). The more in-depth genetic diversity was revealed by Network-Based single nucleotide polymorphism (SNP) discovery protocol. It suggested the differentiation of switchgrass groups by isolation-by-ploidy, the migration from south to north, and the incidence of tetraploid upland from octaploid upland (Lu et al., 2013). Such a large diversity of switchgrass populations can provide a source for disease resistance (Hoisington et al., 1999).

Plant-pathogen interactions are complicated. The basal defense is initiated when conserved molecular signatures of pathogens called pathogen-associated molecular patterns (PAMPs) are recognized by plant pattern recognition receptors (PRRs). Such a recognition activate PAMP-triggered immunity (PTI). Successful pathogens can suppress PTI by secreting virulent effector proteins. These effectors then trigger the second line of defense called effector-triggered immunity (ETI) mediated by resistance R genes (Jones & Dangl, 2006). In biothrophic pathogens, a gene-for-gene interaction between resistance R genes of the plant and avirulent (Avr) genes of the pathogen result in a hypersensitive response leading to local programmed cell death and and the end of colonization, confering resistance. Since *B. oryzae* is a necrotrophic pathogen (Ou, 1985), the interaction between switchgrass and the fungus was modelled potentially as an inverse gene-for-gene model (Friesen et al., 2007). In general, the pathogen secretes necrotrophic effectors as host-selective toxins (HST), such as SnTox1 from *Stagonospora nodorum*, T-toxin from *Cochliobolus heterostrophus*, and HC-toxin from *C. carbonum* (Liu et al., 2004; Ullstrup, 1972; Walton, 1996), that interact with host sensitivity genes, resulting in a compatible susceptible interaction to trigger host cell death (Friesen et al., 2008). Such an interaction is known as effector-triggered susceptibility (ETS). Although crude extract from *B. oryzae* was proposed to contain HST (Vidhyasekaran et al., 1986), the toxin has never been characterized. Instead, according to comparative genome study (Condon et al., 2013) and screening on a wide range of hosts (Chakrabarti, 2001; De Bruyne et al., 2016), *B. oryzae* does not produce any HST but produces ophiobolin A and B as a non-host-selective toxin (Xue et al., 2015) triggering many pathways such as reactive oxygen species (ROS) detoxification, protein phosphorylation, and ethylene production (Kim et al., 2014; Bockhaven et al., 2015).

Such an ETS interaction suggested that susceptible cultivars can be rapidly screened for pathogens carrying the necrotrophic effectors (Juliana et al., 2018). Breeding for the improvement of resistance to BLS can be done by eliminating susceptible alleles from the population. However, the recurrent phenotypic selection for the resistance in upland switchgrass cultivars ‘Shelter’ and ‘Cave-in-Rock’ for two cycles of selection did not provide any more resistance to BLS (Songsomboon, 2019). The screening was done in a seedling stage; therefore, future breeding for the resistance with various screening methods, in various growth stages, and bigger plant populations was suggested. Identification of resistant genotypes or populations is still needed.

To accelerate breeding for disease resistance, genomics-assisted breeding is an important approach (Leng et al., 2017). Basically, it begins with gene identification, isolation, cloning, functional characterization, validation and utilization. There are two main approaches to identify resistance genes in a diverse population: linkage mapping and genome-wide association studies (GWAS). Although GWAS cannot confirm the causal polymorphism, it depends on linkage disequilibrium (LD) that potentially link between the markers and the causal polymorphisms. The diversity panel in GWAS contains high allelic diversity and ancestral recombination events resulting in a finer resolution than linkage mapping (Yu and Buckler, 2006). The technique has been used to dissect flowering time in switchgrass (Grabowski et al., 2017). Although GWAS has never been used to dissect disease resistance in switchgrass, it has successfully dissected resistance to leaf rust caused by *Puccinia triticina* Eriks., tan spot caused by *Pyrenophora tritici-repentis* (Die.) Shoemaker and stripe rust caused by *Puccinia striiformis* West. in wheat (*Triticum aestivum* L.) (Juliana et al., 2018). Despite no resistance genes identified in switchgrass, there were 13 SNP markers from quantitative trait loci (QTLs) linked with the resistance in rice (Sato et al., 2008, 2015). This study will provide the first dissection of the resistance to BLS in switchgrass.

The objectives were 1) to determine genotypes or populations from the northern switchgrass association panel that can be candidates for resistance to BLS, 2) to conduct a single- and multi-trait GWAS for resistance to BLS in the association panel via both natural and artificial inoculation, and 3) to explore the genes linked to the significant markers from the GWAS model to determine potential candidate genes for resistance to BLS.

## MATERIALS AND METHODS

### Switchgrass in the Northern Association Panel

The switchgrass population used in this study had been developed for accelerating breeding progress especially for bioenergy traits at northern latitudes (Lu et al., 2013; Lipka et al., 2014). The association panel consists of 478 plant genotypes from 66 populations representing mostly the upland northern population and some southern lowland populations. ‘Population’ in this study means a seed source with a specific origin and, in some cases, breeding history. Six to ten seeds from each population were planted in to individual plants called “genotypes”. The association panel was initially planted in Ithaca, NY, in 2008 and then vegetatively cloned and planted in a randomized, complete block design with three replicates in 0.9 m^2^ spaced planting in Ithaca, NY, and Philipsburg, PA, in 2016. No fungicide had been applied to the field.

### Disease evaluation and phenotype processing

Since the unsuccessful recurrent phenotypic selection for resistance to BLS was done in the seedling stage (Songsomboon, 2019), the phenotypic evaluations in this study were explored at a mature stage and different conditions. Bipolaris leaf spot has been evaluated in both fields to determine the resistance from natural incidence under different environments and laboratory to minimize environmental effects on the resistance. Field evaluation was conducted in the same week in both locations in August 2017. Each plant was visually evaluated by one person to reduce the variance among persons. The disease severity score ranged from 0 to 5, as 0 = 0% BLS, 1= 1-10% BLS, 2= 11-25% BLS, 3= 26-50% 134 BLS, 4= 51-75% BLS, and 5= 76-100% of the leaf area of the whole plant with BLS. In addition to field evaluation, to minimize environmental effects, an artificial inoculation in the laboratory was conducted. Disease severity was evaluated by detached leaf and leaf disk assays by taking leaf samples from one replicate of genotypes in Ithaca, NY. In the detached leaf assay, three healthy leaves were randomly selected from 30 centimeters below the top of the shoot to control the same stage of leaf development. The leaves were cut into five centimeter segments, washed with deionized water, placed in a pre-wet petri dish bedded with filter paper, and kept cold in a cooler in the field, then refrigerated overnight.

The inoculum was prepared from a subculture of a single conidium isolated from a ‘Carthage’ switchgrass leaf in the warm season biofuels field experiment in Ithaca, NY (Waxman, 2011). The day after collecting the detached leaf, the three-week-old plate was flooded, filtered via gauze, and the concentration adjusted to 10^5^ conidia×ml^−1^ by hemocytometer with 2 drops of Tween-20 per 100 ml. An airbrush pressuring at 10 psi was used to spray inoculum on each dry-surfaced adaxial side of the leaf in each plate. After letting the droplets of inoculum dry and firmly attach to the leaf surface, the plates were then sealed with a paraffin tape to maintain high moisture condition. Each plate contained three detached leaves from one genotype. The plates were placed on table randomly and kept at room temperature with 12-hour light for seven days showing necrotic lesions and leaf necrosis (Figure 1). Due to the limitation of laboratory space for the plates and labor in the leaf collection, 85 to 200 genotypes were sampled in each set of the experiment, with some overlapping genotypes in one to three duplicated plates and additional non-inoculated plates as controls. The leaf samplings were done weekly from May to June 2017 but were controlled by the attempt to sample the similar stage of leaf by sampling from the same height from the top. A total of seven sets of the experiment eventually covered all of 478 genotypes. The disease evaluation was conducted by vision and by image analysis software. Both evaluations were based on percentage of leaf covered with lesions. The evaluation by vision was from 0 to 100 % by 10% increments. Then, each leaf was dried and scanned with a flatbed scanner (Canon CanoScan LiDE 700F) at a resolution of 1,200 dpi, and images were saved in .tiff format. Although the detached leaf assay was an unbalanced design, best linear unbiased predictions (BLUPs) to extract random effect of the genotypes was suggested to be able to alleviate the problem by fitting the experiment factor as a fixed effect (Hill & Rosenberger, 1985; Piepho et al., 2008; Robinson, 1991; Stroup & Mulitze 1991; White & Hodge, 2013).

**Figure 1.**
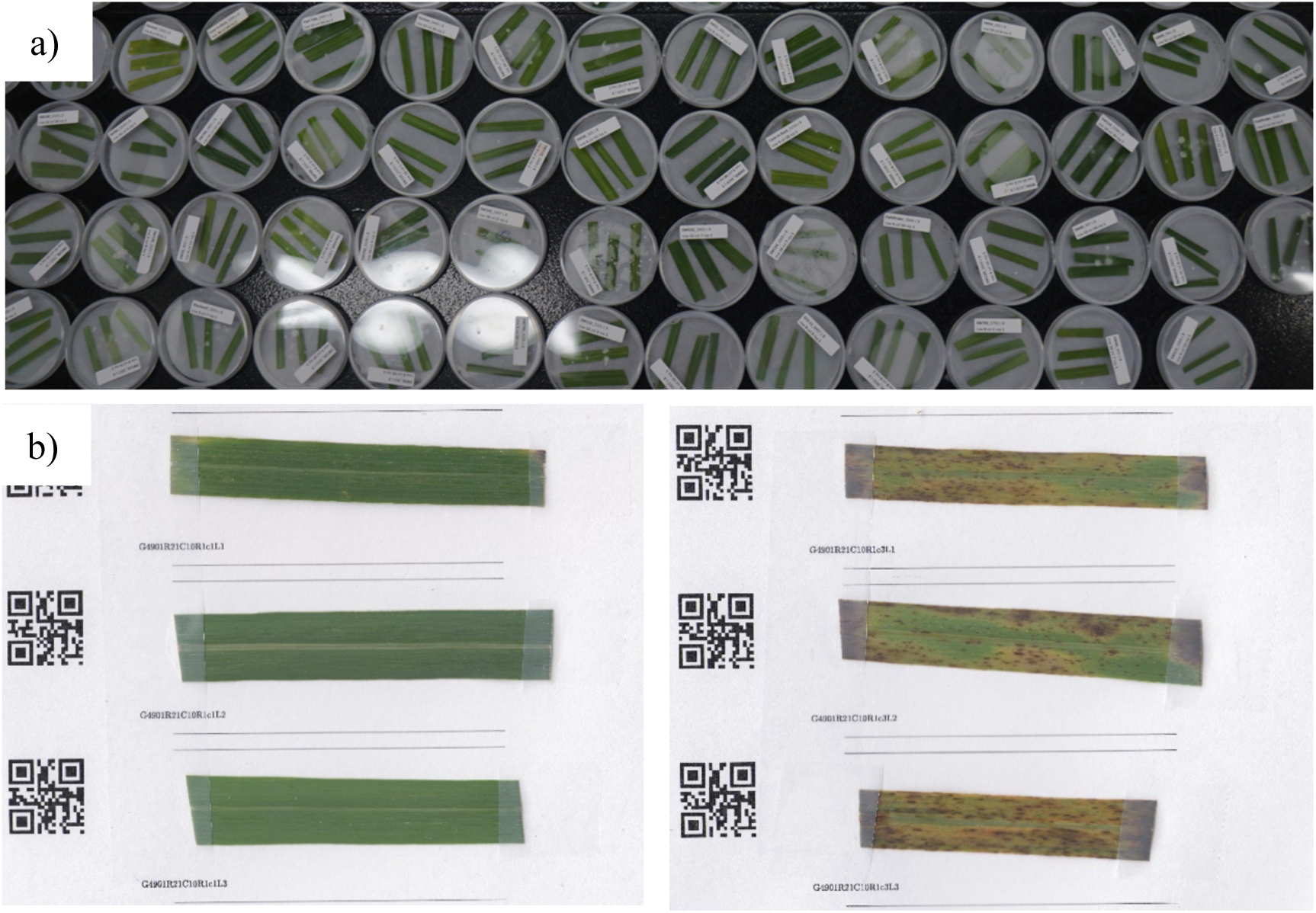
Detached leaf assay; a) sealed plates of detached leaves under 12-h light 25°C; b) control inoculated (left) and inoculated leaves (right) at 7 dpi were taped on white paper with QR code label for scanning for image analysis.

In the leaf disk evaluation, all of 478 genotypes were collected on the same day in August 2017 by cutting the same age leaves from each genotype and keeping them refrigerated overnight. On the next day, the leaves were washed with deionized water, surface dried, bored by keeping the midrib in the middle into an 8-mm disk, and placed on a water agar plate. There were 28 leaf disks in each plate, including four check disks (non-inoculated control and inoculated disks of resistant ‘SW43_09’ and susceptible ‘SW122_02’) and three replicated disks of eight genotypes (Figure 2). The inoculum was prepared the same way described above. For each leaf, two µL of inoculum was dropped in the middle of the right side of the leaf. After letting the droplet dry, the total of 64 plates were sealed with paraffin tape to maintain high moisture, randomly placed on a table, and kept under 12-hour light at room temperature. The disease evaluation was conducted at seven dpi both by vision and image analysis software. The percent of leaf disk covered with lesions and necrotic tissue was assessed by vision from 0% to 100% with 10% increments. Also, on the same day, a photo was taken of each plate with a digital DSLR Canon Rebel T6 DSLR camera with the resolution of 2644 x 3084 in .JPG.

**Figure 2.**
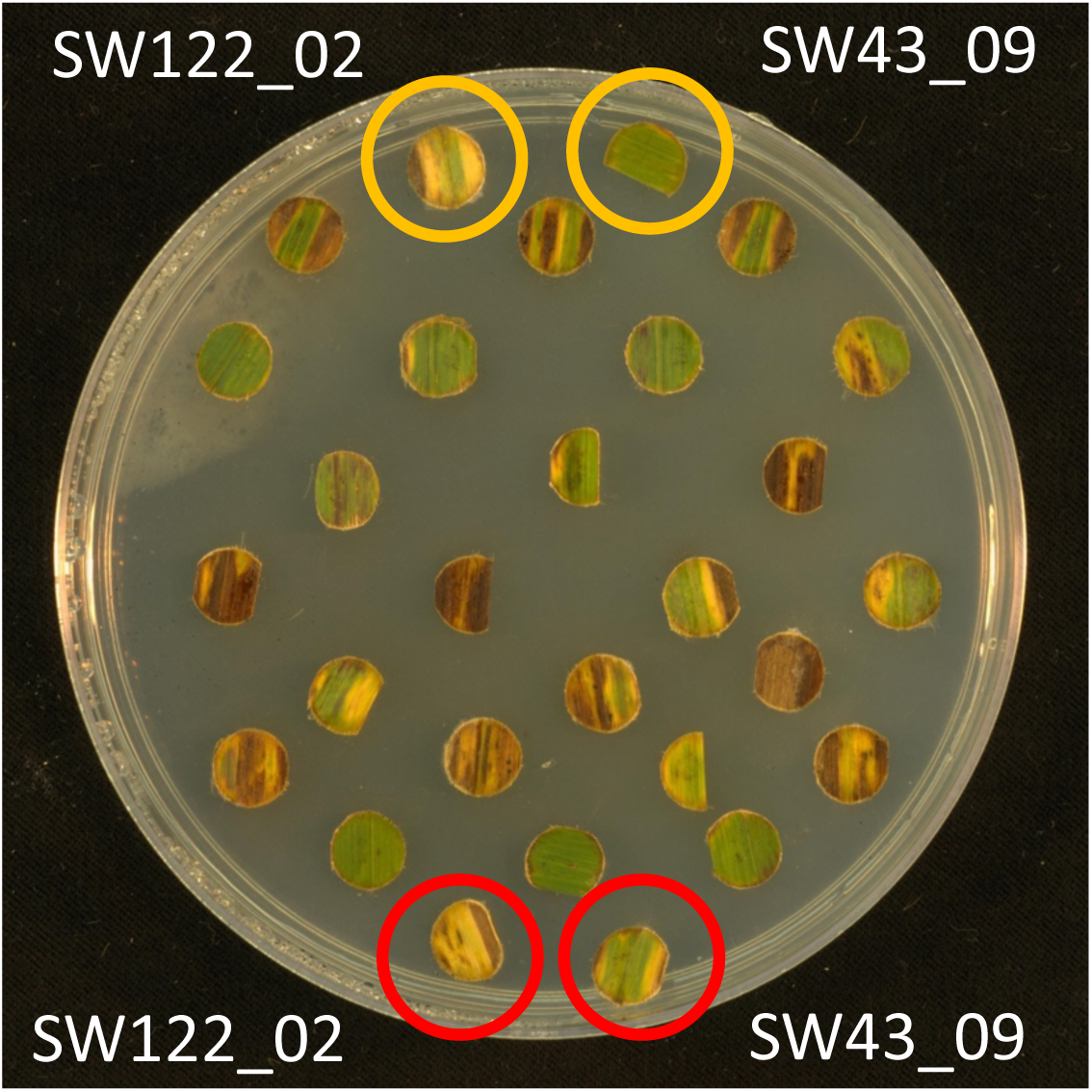
A plate example of leaf disk assay at 7 dpi. Each plate consisted of 28 leaf disks. SW43_09 was used as a resistant check and SW122_02 as a susceptible check. Two non-inoculated control leaf disks (yellow circles) and two inoculated leaf disks (red circles) were included in all plates. The rest of the leaf disks (24) consisted of eight genotypes with three replicates placed in a consecutive row from left to right.

Images from both detached leaf and leaf disk assays were analyzed by using ImageJ macro at the setting provided in the Supplement file. In brief, each detached leaf or leaf disk was measured for total leaf area and total necrosis lesion area (yellow and black area). Percentage of area covered with lesions was computed by dividing the total necrotic lesion area by total leaf area.

Package ‘LME4’ (Bates et al., 2015) in R was used to calculate BLUPs from various disease evaluations. For the field evaluation, BLUPs were computed based on each location separately and on the two locations combined using genotypes nested in populations and replicates as random effects in each location model and populations as a fixed effect. BLUPs for the two locations combined were fitted as genotype nested in population, replicates, locations, and the interaction between genotypes and location as random effects and populations as a fixed effect. In addition to generating BLUPs from the field evaluations, the highest score, which is the most severe symptoms observed from each location and the combined two locations over replicates, were used based on the successful example of QTLs of resistance to foliar symptoms caused by potato virus Y in autotetraploid potato (*Solanum tuberosum* L.) (Silva et al., 2017). The highest scores were used on the assumption that in the field there was a chance that some replicates may be exposed by or avoid the pathogens differently. The maximum scores were considered the worst response of each genotype. For disease evaluations under laboratory condition, since all phenotypes were expressed as percentages to leaf area, log transformation was performed before computing BLUPs. For the detached leaf assay, BLUPs were computed with experiment, genotype nested in populations, interaction between genotypes and experiment, plates within the experiment, and leaf replicates of genotypes as random effects and populations as a fixed effect. For the leaf disk assay, BLUPs were calculated with plates as a fixed effect and genotype, replicates within plates, and genotype within plates as random effects and populations as a fixed effect. Broad-sense heritability was estimated from variances in each model and standard error was computed by bootstrap approach for 1000 times (Supplement file). Moreover, phenotypic correlations were computed between resistance to Bipolaris leaf spot to the 20 morphological and biomass quality traits from Lipka et al. (2014), such as plant height, anthesis date, acid detergent lignin, minerals, ethanol·g^−1^ dry weight, etc.

Therefore, the resistance to BLS was evaluated via three approaches – detached leaf, leaf disk, and field evaluation – providing phenotyped traits including BLUPs of detached leaf percent lesion via vision and image analysis (DTVI and DTIA), BLUPs of leaf disk percent lesion via vision and image analysis (DSVI and DSIA), BLUPs of field evaluation from two locations, NY and PA (BTL, BNY, and BPA), highest score from two locations, NY and PA (MTL, MNY, and MPA).

### Genotyping, linkage disequilibrium analysis, and population structure

HAPMAP v.1 set from Evans et al. (2015) provided 1,377,841single-nucleotide polymorphisms covering 38,654 genic regions. In short, DNA of each genotype was processed via exome-capture using the Roch-Nimblegen switchgrass exome-capture probe set (Evans et al., 2014) and DNA sequencing. The sequences were aligned to the *P. virgatum* genome assembly v.4.1 (*P. virgatum* v.4.1, DOE-JGI, http://phytozome.jgi.doe.gov/) for SNP discovery (Ramstein et al., 2018). The reads were sorted with PicardTools version 2.1.1 (http://broadinstitute.github.io/picard) SortSam and MarkDuplicates. Samtools version 1.3.1 (Li et al., 2009) was used to pile up files with base alignment quality disabled and map quality adjustment disabled. In this calling, SNPs were required to be bi-allelic, sequenced in all samples, be monomorphic in at least two samples, filtered a minimum read mapping quality score of 30, and more than 5X coverage in more than 95% of the samples. At each SNP, the genotype dosages ranged from zero to two copies of minor alleles and can be nonintegers by using EM algorithm (Martin et al., 2010). Naturally, the switchgrass association panel includes allopolyploid switchgrass: tetraploids (4x), octoploids (8x) and hexaploids (6x). It is challenging to perform GWAS with polyploid models. In this study, we modelled them under the assumption of disomic inheritance similar to the study of GWAS of flowering time in Grabowski et al. (2017). The reason was that tetraploid switchgrass was confirmed to show disomic inheritance (Okada et al., 2010). Although octaploid switchgrass had four copies of each homologous chromosome, which was difficult to precisely determine heterozygous genotypes, the disomic segregation was used for the model with the caution of the increasing standard error of estimates in GWAS. Moreover, linkage disequilibrium (LD) was computed via ‘-r2’ tag in PLINK as r^2^ between all SNPs within 1 MB and recorded all values (Purcell et al., 2007). Principle component analysis was used to evaluate the population structure via PCA in FactoMineR package (Lê & Husson, 2008). Additionally, ADMIXTURE was used to evaluate population structure with five-fold cross-validation (Alexander et al., 2009).

We were also interested in testing whether the variations in phenotypes were from the high differentiation across five ecotypes (lowland north, lowland south, upland east, upland north, upland west) based on Qst-Fst analysis. Estimated Qst-Fsts were computed by R package “QstFstComp” by bootstrapping 1,000 times by sampling random genotypes (Gilbert & Whitlock, 2015).

### Genome-wide association studies

Genome-wide Efficient Mixed Model Association (GEMMA) (Zhou and Stephens, 2012) was used to implement a multivariate linear mixed model for GWAS of single- and multiple-phenotyping approaches for resistance to BLS, to analyze each phenotype and combined phenotypes for improving statistical power (Porter & O’Reilly, 2017; Zhu et al., 2015). Kinship was included as a random effect. Also, the first three principal components (PCs) were included as fixed covariates. Although the results of Bayesian information criterion resulted in no improvement of the GWAS model with PC inclusion (Lipka et al., 2013), the inclusion of PCs helped to improve the QQ-plot closer to the theoretical QQ-plot for this resistance to BLS. The GWAS was conducted in five groups including all of 478, tetraploid (4X), octaploid (8X), lowland, and upland genotypes to determine if there were any SNPs linked to specific switchgrass populations. Minor allele frequency (MAF) at 0.05 was used to filter the minor alleles. In total, 135,791 SNP markers were used in the association analysis. To correct multiple testing, false discovery rate (FDR) was calculated as 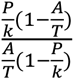 where P is the p-value tested at 0.1, k is number of traits in the combination, A is the number of SNPs that were significant at the p-value tested and T is the total number of SNPs tested. The significant markers then were used to determine candidate genes linked to them within the range of LD and search in JBrowse in *P. virgatum* v.4.1 in Phytozome (e.g., if the position of the marker was 4019500 on chromosome 9K and LD was 20 kb, “Chr09K:4019500..4039500” was used). In case that the candidate gene was not fully aligned within that LD range, we considered the gene as a candidate if 80% of its length was within the LD range. Since some of SNPs’ positions could not be aligned to the *P. virgatum* genome assembly v.4.1, they were excluded from determining the candidate genes.

### Candidate genes based on resistance in rice

Since *B. oryzae* is one of the major pathogens in rice (*Oryza sativa*), major resistance QTLs have been identified in recombinant inbred lines (RILs) and double haploids (DH) by using restriction fragment length polymorphism (RFLP), simple-sequence repeat (SSR) markers, and sequence-tagged site (STS) (Sato et al., 2008, 2015; Katara et al., 2010). Three major QTLs in Chromosomes 1, 4 and 11 were then studied further in near-isogenic lines (NILs) and 13 SNP markers linked with the resistance (Sato et al., 2015). The NILs are BC3F5 between indica ‘Tadukan’ (resistance) and temperate japonica ‘Koshihikari’ (susceptible). According to GWAS in rice, linkage disequilibrium (LD) are ∼ 100 kb in indica and ∼200 kb temperate in japonica (Zhao et al., 2011). Thus, in this study, the candidate resistance genes were screened from 200 kb around the 13 SNPs from *Oryza sativa* genome assembly v.7.0 (*Oryza sativa* v.7.0, DOE-JGI, http://phytozome.jgi.doe.gov/). The candidate genes from rice are listed in the table S1. The sequences of these candidate genes were used to identify potential homologs in switchgrass by using the BLASTP tool on Phytozome and kept the hits with top 10% highest score (Goodstein et al., 2012).

## RESULTS

### Phenotyping and correlation between traits

The results from detached leaf showed that ‘ECS.6’ had the lowest severity (DTVI 24% and DTIA 8%) and ‘SW115’ had the highest severity among populations (DTVI 81% and DTIA 60%) (Table S2). However, they were not significantly different from the other population with lowest and highest severities (Table Stat1&2). When comparing among genotypes, ‘SW788.05’ had the lowest severity (DTVI 6% and DTIA 2%) whereas ‘SW63.05’ had the highest severity (DTVI 100% and DTIA 100%) (Table S3). Similarly, they were not significantly different from the other genotypes with lowest and highest severities (Table Stat11&12). The leaf disk assay showed that ‘SW803’ had the lowest severity (DSVI 20% and DSIA 48%) and ‘SW38’ had the highest severity (DSVI 98% and DSIA 99%) but not significantly different than other populations (Table Stat3&4). In the genotype-based comparison, ‘High Tide.02’ had the lowest severity (DSVI 10% and DSIA 11%) but not significantly different than other genotypes (Table Stat13&14), and the highest severity at DSVI 100% and DSIA 100% among the 81 genotypes. Based on artificial inoculations, both the detached leaf and the leaf disk assay suggested that the lowland ecotypes (DTVI 43%, DTIA 21%, DSVI 43% and DSIA 70%) appeared to be more resistant than the upland ecotypes (DTVI 61%, DTIA 39%, DSVI 67% and DSIA 89%) which are statistically significant (p-value < 0.05) (Table S4).

In the field evaluation based on the mean of the severity score (0 to 5) from two locations, ‘Shelter’ showed the lowest severity (0.44) and ‘SW787’ the highest severity but not significantly different from the other populations (Table Stat5). However, when considering each location, different populations performed differently. In comparison between NY and PA, Wilcoxson’s rank test showed significant difference between the two locations (p-value=0.042). The re-ranking incidence (Le et al., 2001; Muir et al., 1992) across locations indicates the significant GxE effect. In NY, SW803 had the lowest severity (0.22) and SW787 had the highest severity (3.0) despite not significantly different from other populations (Table Stat6). Whereas, in PA, Shelter had the lowest severity (0.33) and Pathfinder had the highest severity (2.9) despite not significantly different from other populations (Table Stat9). Between the two locations, ‘SW123’, ‘SW33’, ‘SW793’, ‘SW781’, High Tide, and Timber had the lowest MTL at 3 (resistant), and 41 populations had the highest MTL at 5 (susceptible) even being not significantly different from the other 66 populations (Table Stat6). In NY, SW123, SW128 and ‘ECS.6’ had the lowest MNY at 2, and 32 populations had the highest MNY at 5. In PA, ‘SW115’, ‘SW802’, ‘SW31’ and ‘SW43’ had the lowest MPA at 2, and 19 populations had the highest MPA at 5 without significant differences (Table Stat8&10). Although, each phenotyping approach cannot determine single resistant population, a combination of detached leaf, leaf disk, and mean from two locations showed that ‘SW788’, ‘SW806’, ‘SW802’, ‘SW793’, ‘SW781’, ‘SW797’, ‘SW798’, ‘SW803’, ‘SW795’, ‘SW805’ were always in the lowest severity group and potential candidates of resistance.

When comparing among ecotypes, the field evaluation of means between two locations showed the trend of more resistance in lowland than upland ecotypes significantly (Figure S1 & Table S4). The same difference also occurred in NY and PA. The severities based on genotypes in each phenotyping approach are listed in Table S3 and Stat15-18.

Broad-sense heritabilities (H^2^) in different groups of genotypes were similar across groups with some variations (Table 1). For example, H^2^ from DSIA in upland was only 0.14 while other groups had around 0.40. Although each trait from different phenotyping approaches in different groups had significantly moderate to high H^2^, H^2^ of severity from the two locations combined was zero. This was supported by the significantly different mean of severity from the two locations, NY and PA (Figure S2 and Wilcoxson p-value = 0.042). Due to zero heritability, the BTL was zero and cannot be used for conducting GWAS. The different ranking was not only present in the field evaluation, the other phenotyping approaches yielded different ranking of resistance as suggested by the ranks of DTIA, DSIA and mean from two locations and the correlations between the traits (Figure S2, Figure 3 and Wilcoxson’s p-value<2.2e-16). For example, from 478 genotypes, ‘KY1625_07’ ranked 27 from DTIA but ranked 329 in DSIA, and 100 in mean from two locations. Such differences led to low correlation among approaches such that the r^2^ between DTIA and DSIA was only 0.1 and between DTIA and mean of two locations (TL) was 0.03 (Figure 3). In contrast to these low correlations, there were high correlations in DTVI-DTIA (r^2^ = 0.84) and DSVI-DSIA (r^2^ = 0.85). There was no BLS resistance trait correlating to other agronomic or biomass quality traits (Figure S3).

**Figure 3.**
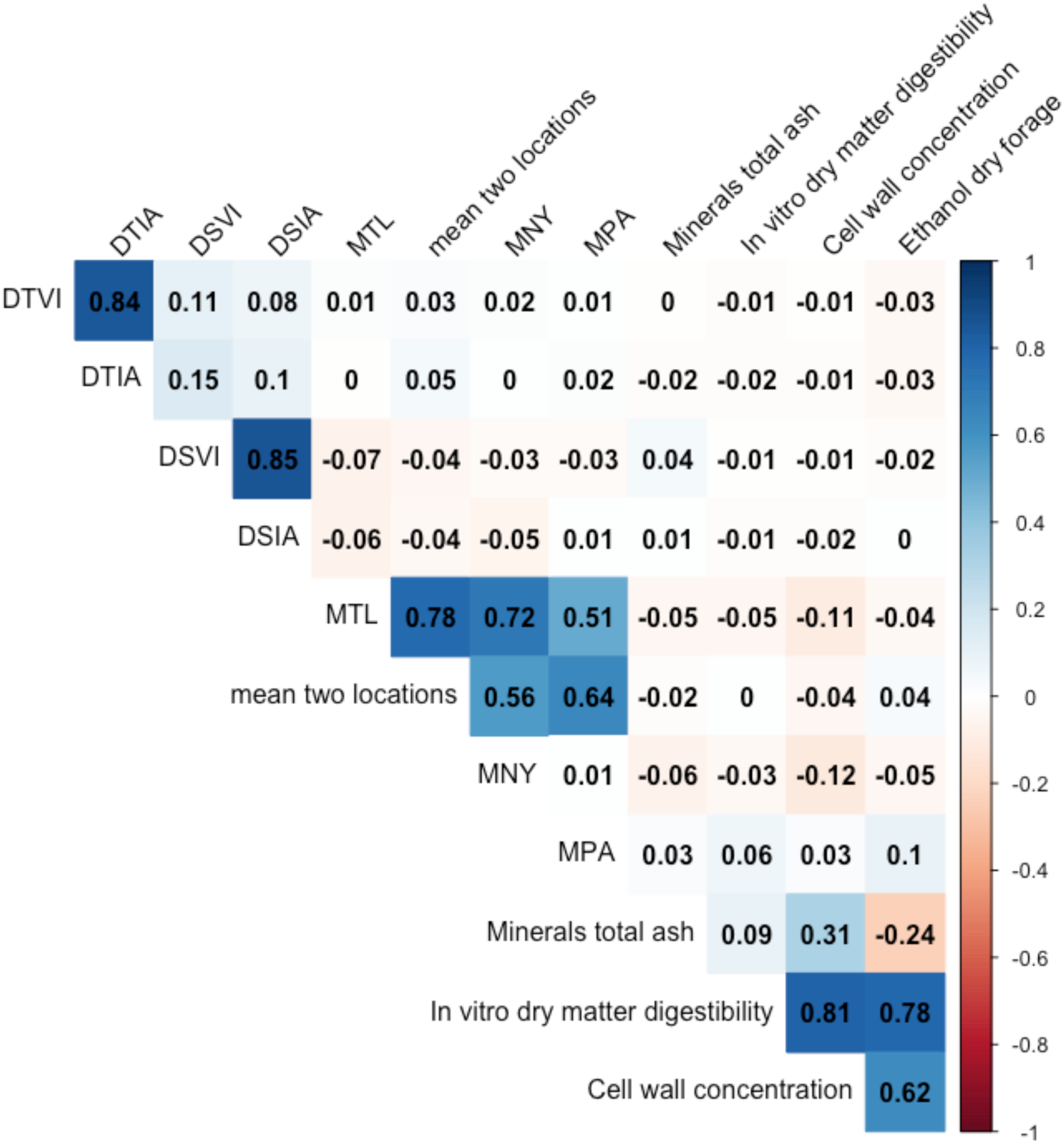
Correlation plot among BLUPs of severity from the detached leaf via vison (DTVI), via image analysis (DTIA), from leaf disk assay via vision (DSVI), via image analysis (DSIA), the highest score between two locations (MTL), mean from two location (TL), minerals total ash, *in vitro* dry matter digestibility, cell wall concentration, and ethanol conversion per day forage.

**Table 1.**
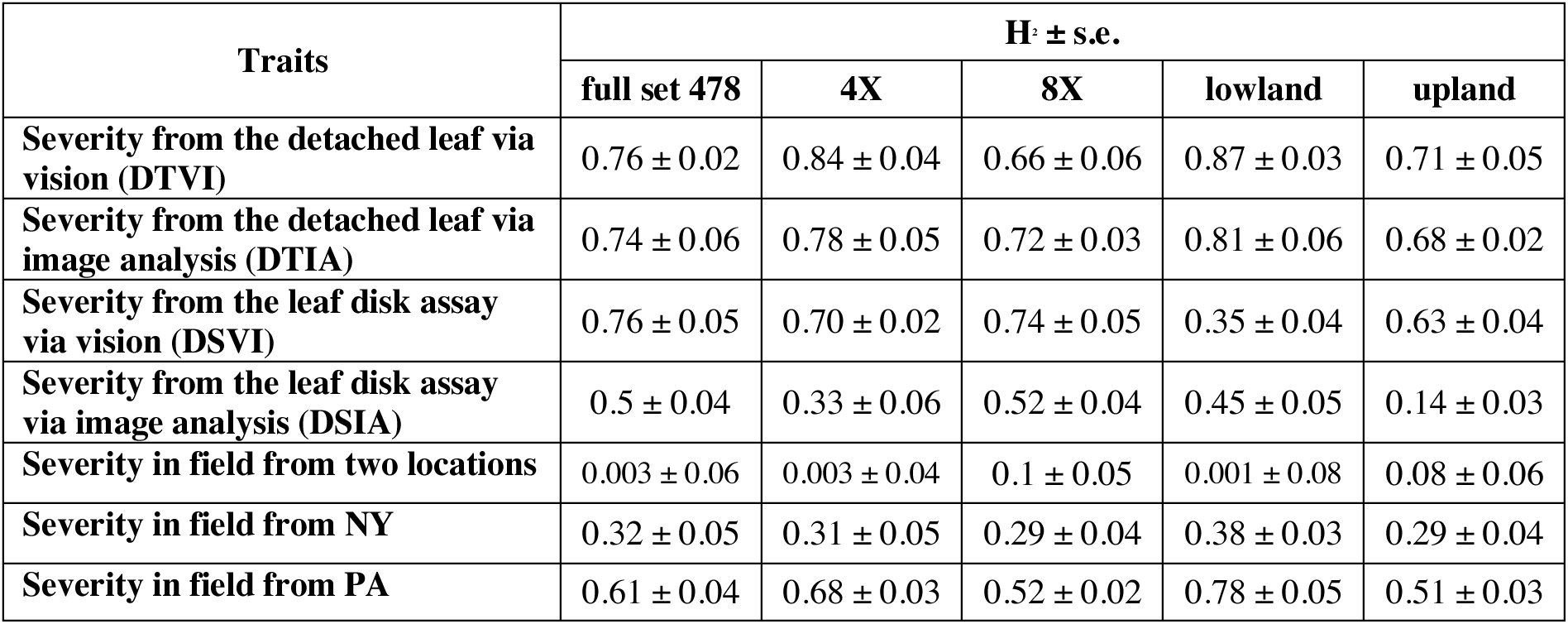
Broad-sense heritability of each trait in each groups.

In addition to comparing solely on phenotype, Qst-Fst comparisons were used to investigate whether the mean difference of each of the phenotypes among populations is greater than expected mean difference under genetic drift. None of the traits (DTVI, DTIA, DSVI, and DSIA) has Qst-Fst significantly more than zero (Figure 4 and Figure S4) showing no positive selection for resistance to BLS in populations.

**Figure 4.**
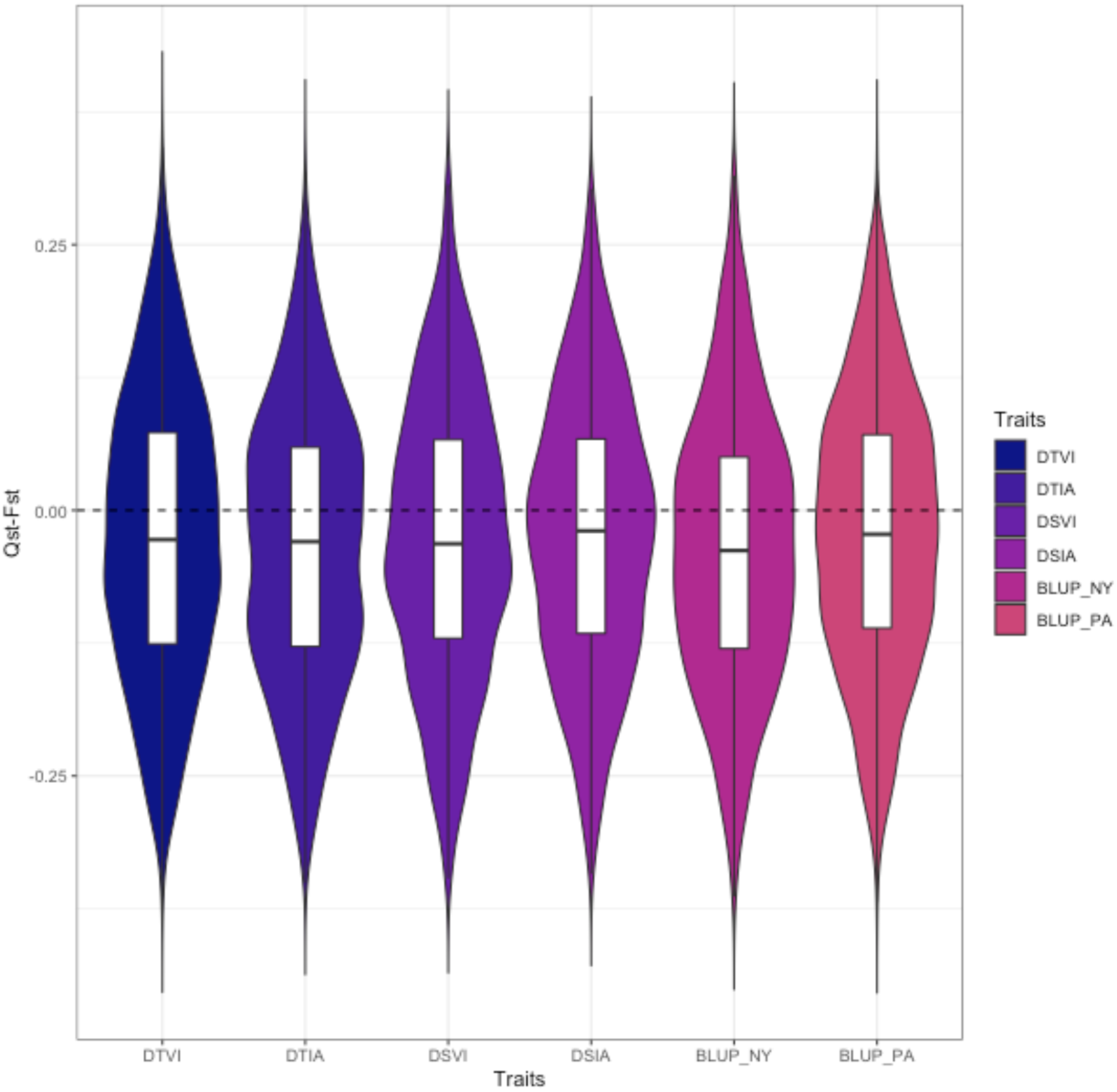
The bootstrapped (1000 times) distributions of Qst-Fst for each phenotype (DTVI, DTIA, DSVI, DSIA, BLUPs NY and BLUP PA) are compared to the expected value of zero under neutrality (black dashed line).

### Linkage disequilibrium, principal component analysis and admixture

To determine the interval that potentially linked to the significant markers from GWAS, linkage disequilibrium (LD) decay was estimated by plotting the allele frequency correlations (R^2^) against the physical distance in base pairs (Figure 5). In the full set of 478 genotypes, the LD decayed sharply within 20 kb and reached a plateau background within 50 kb. Therefore, in this study, we focused on genes within the 20 kb from the significant SNPs.

**Figure 5.**
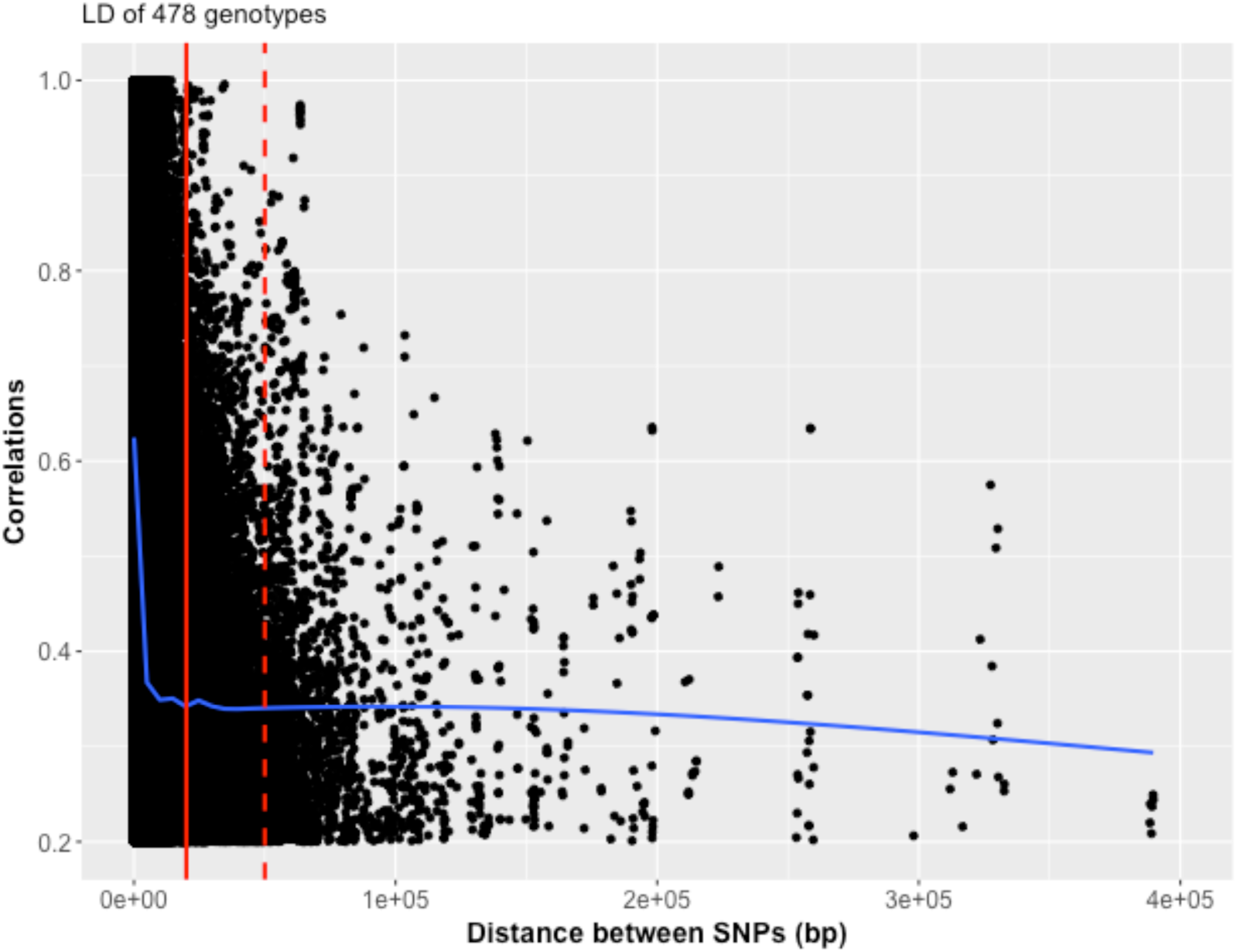
Scatter plot showing the linkage disequilibrium (LD) decay by plotting physical distance in base pairs against the LD estimate as correlations (R^2^) in 478 genotypes. The LD (blue curve) decays rapidly within 20 kb (red line) and reaches background levels around 50 kb (dashed red line).

Based on the principal component analysis and admixture, there was a population structure in this association panel (Figure 6). From PC1 and PC2, upland and lowland ecotypes separated from each group. Also, within the lowland ecotype, the latitude of lowland can be distinguished to lowland north and south groups. The separation of upland ecotype can also be noticed in PC1 and PC3 in that the upland north was grouped apart from the other upland. Besides ecotypes, ploidy level can be differentiated into groups. Both lowland north and south, and upland north genotypes were tetraploid (4X) while the upland east and upland west genotypes were octaploid (8X). In total, PC1, PC2, and PC3 explained the variance due to the population of 49.35%. The admixture (Figure 5b) also showed the distinction among five groups of ecotypes.

**Figure 6.**
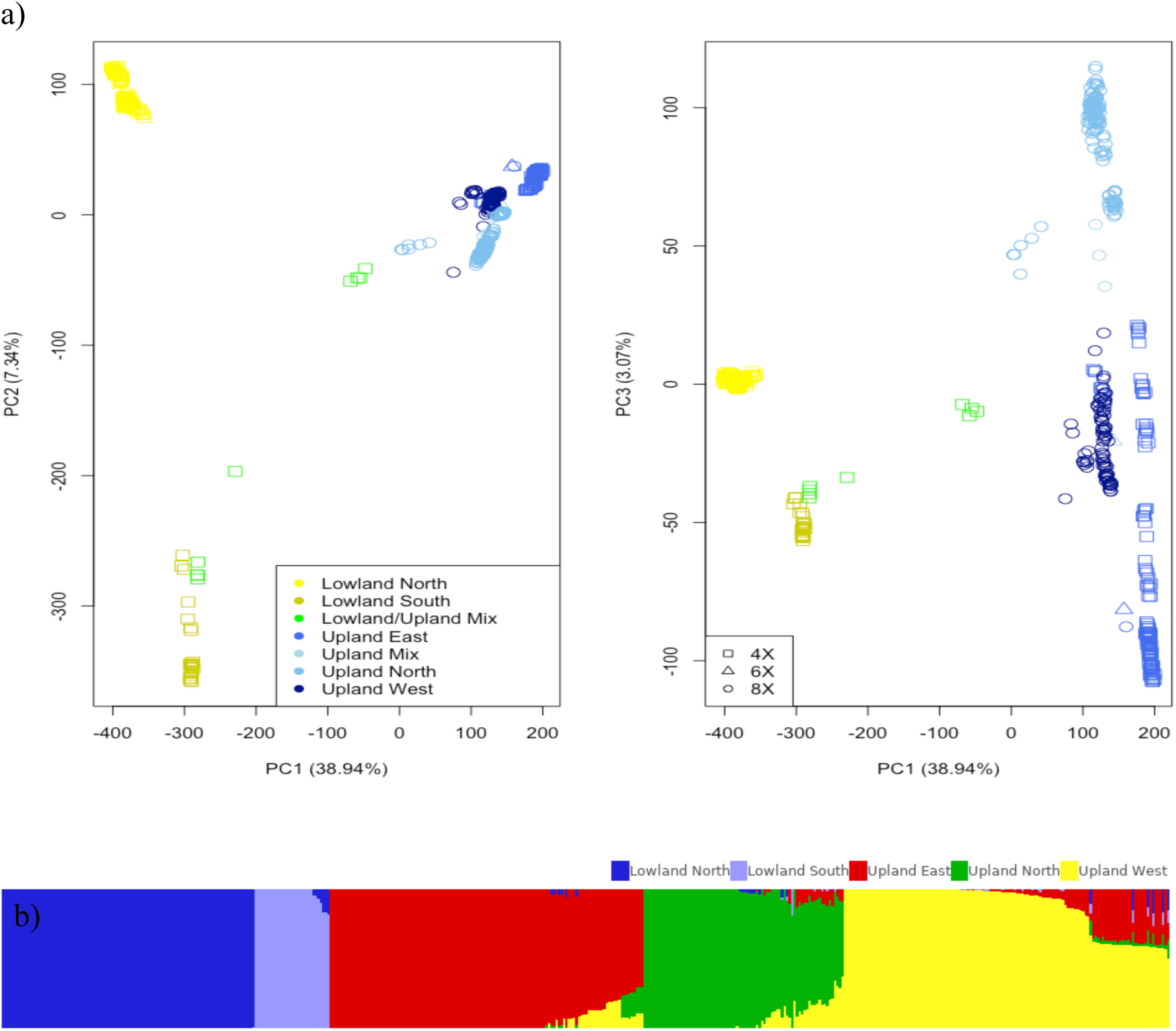
a) The 478 switchgrass genotypes from the Northern Association Panel showing the different distributions based on ecotype and region from principal component analysis (PC). Colors represented different ecotypes and shapes represented different ploidy levels. Percentage variances explained by PCs are in parentheses. The lowland ecotype can be differentiated into two regions from PC1 and PC2, and the upland ecotype can be grouped into three regions from PC1 and PC3. b) Admixture also showed the separations of the genotypes into five gene pools.

### Genome-wide association

As we conducted GWAS with five subsets of genotypes including478, 4X, 8X, low- land and upland, most of the single phenotypes did not show significant Manhattan peaks or theoretically-expected QQ-plots (Figure S5 to S8). For multiple combinations of assessing resistance to BLS via many phenotyping approaches, almost all the combinations for GWAS did not provide significant peaks or QQ-plots (Supplement file). To improve the association analysis, the GWASs were then conducted with the combination of three traits of DTVI, DTIA and DSIA, which yielded 14 significant peaks passing FDR threshold and the QQ-plot with the expected distribution (Figure 7a and Table 2).

**Figure 7.**
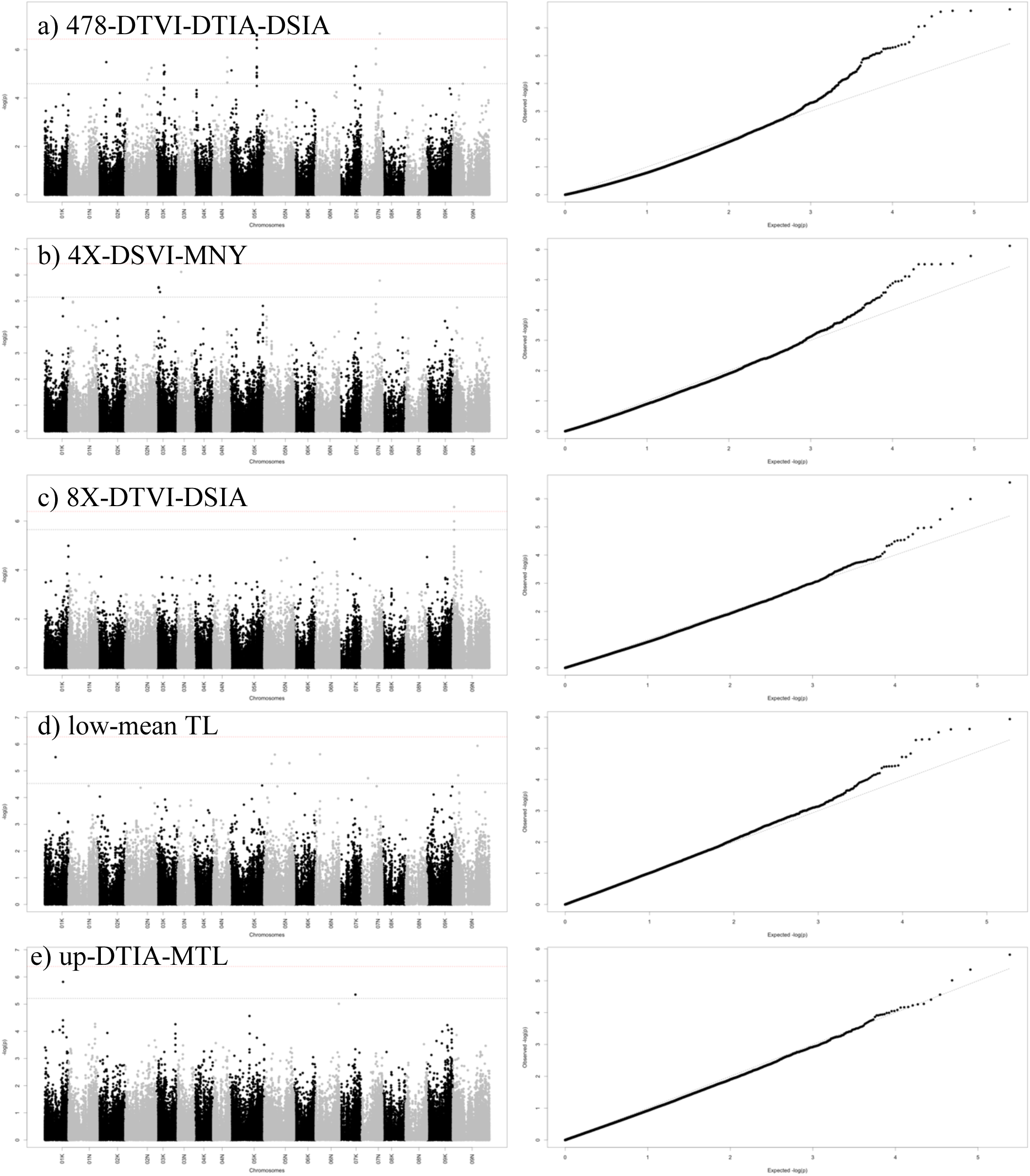
(Left) Manhattan plot showing genetic associated of a) DTVI-DTIA-DSIA in 478 genotypes, b) DSVI-MNY in 4X genotypes, c) DTVI-DSIA in 8X genotypes, d) mean TL in lowland ecotype, e) DTIA-MTL in upland genotypes. The black dashed line represents the FDR threshold (0.1) and the red dashed line represents Bonferroni correction threshold. On x-axis, the physical positions of the SNPs were aligned in 18 chromosomes of *P. virgatum.* (Right) Quantile-quantile (QQ) plots between the distributions of observed to expected P-values for GWAS of each trait combination in each genotype group. Abbreviation: Abbreviation: BLUPs of severity from the detached leaf via vision (DTVI), BLUPs of severity from the detached leaf via image analysis (DTIA), BLUPs of severity from the leaf disk assay via vision (DSVI), BLUPs of severity from the leaf disk assay via image analysis (DSIA), mean of field evaluation of BLS in two locations (mean TL), highest score of BLS in NY (MNY), and highest score of BLS in two locations (MTL).

**Table 2.**
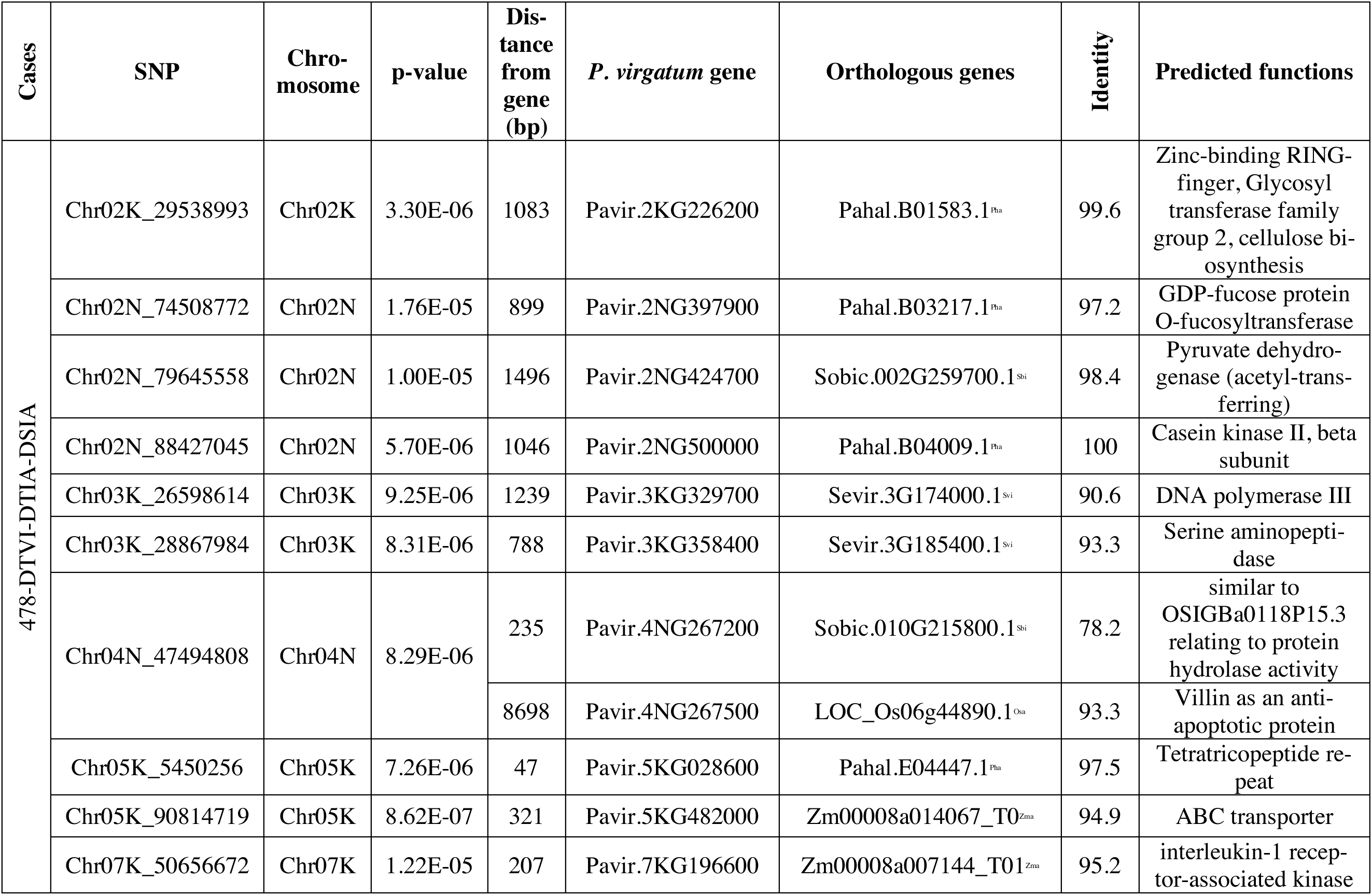
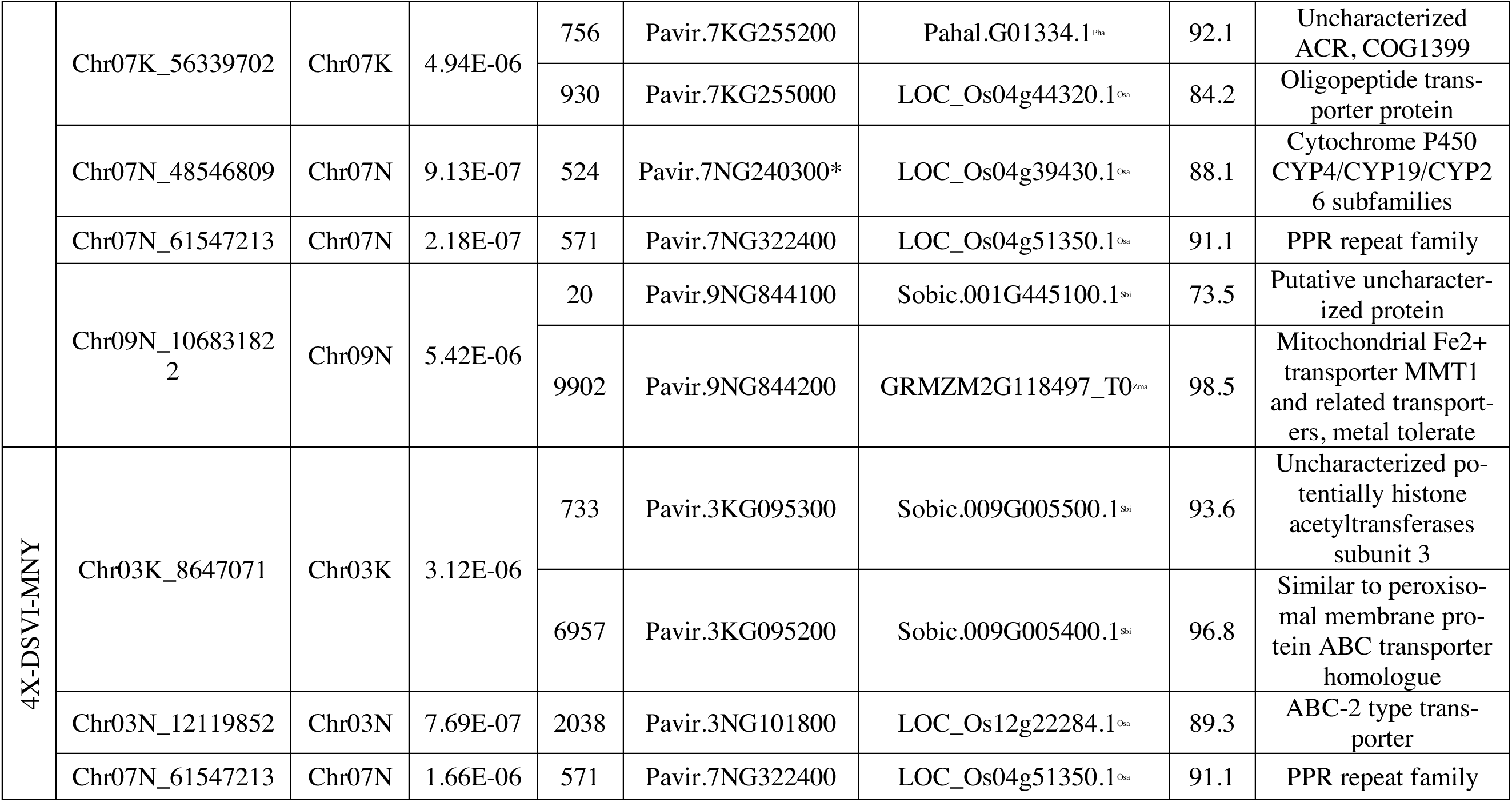
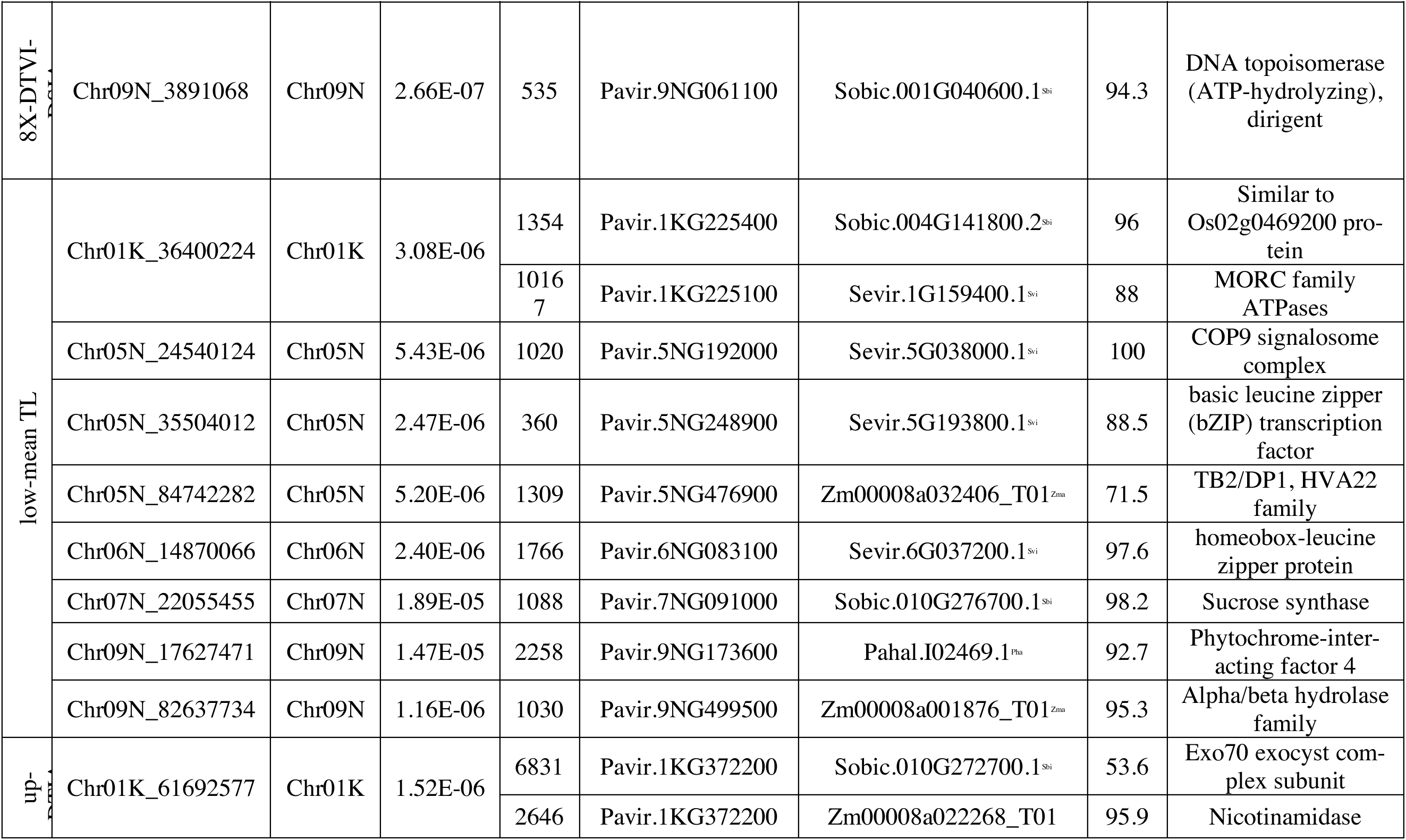
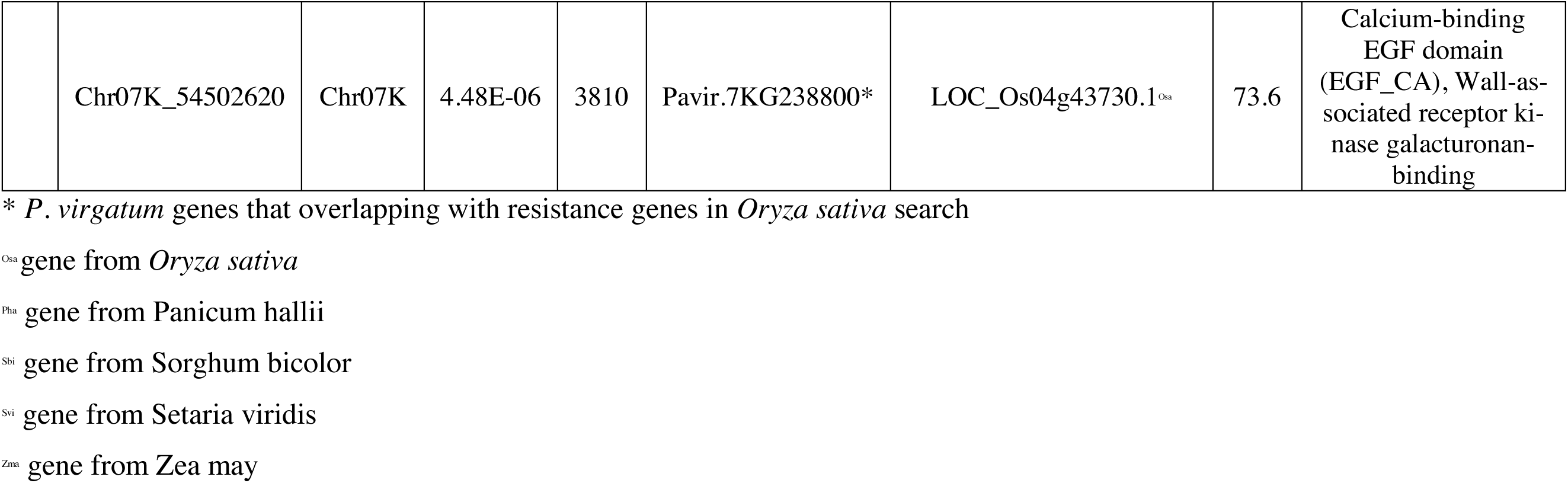
Markers significantly associated with resistance to Bipolaris leaf spot in each case.

In a tetraploid group, the two-trait combination of DSVI and MNY provided three significant peaks with expected QQ-plot (Figure 7b). Interestingly, one peak of SNP Chr07N_61547213 on chromosome 7N from the tetraploid group overlapped with the same peak in GWAS from the 478 group. In the octaploid group, the two-trait combination of DTVI-DSIA showed one significant peak on chromosome 9N (Figure 7c). In the lowland group, the best GWAS was from the single trait mean of the combined two locations yielding eight peaks (Figure 7d). In the upland group, the combination of DTIA-MTL gave two most significant peaks on chromosome 1k and 7K (Figure 7e).

In this study, therefore, we focused on five selected GWAS analyses including 478-DTVI-DTIA-DSIA, 4X-DSVI-MNY, 8X-DTVI-DSIA, low-mean TL, and up-DTIA-MTL (Figure 7). In total, there were 27 significant peaks (one overlap between 478 and lowland group) across 12 chromosomes: 1K, 2K, 2N, 3K, 3N, 4N, 5K, 5N, 6N, 7K, 7N, and 9N. Accumulatively, when considering phenotypic variance explained (PVE) by all 27 significant SNP markers for each trait, PVE of DSIA was the highest at 26.52% and PVE of DTIA was the lowest at 3.28% (Table 3).

**Table 3.** Phenotypic variance explained (PVE) (adjusted PVE) by accumulative 27 significant SNP markers across all analyses for each trait.

| **Traits** | **PVE (adjusted PVE)** |
| --- | --- |
| DTVI | 11.68 (6.38) |
| DTIA | 8.76 (3.28) |
| DSVI | 25.03 (20.53) |
| DSIA | 30.68 (26.52) |
| mean TL | 15.73 (10.67) |
| MNY | 15.74 (10.68) |
| MTL | 13.7 (8.52) |

Abbreviation: BLUPs of severity from the detached leaf via vision (DTVI), BLUPs of sever- ity from the detached leaf via image analysis (DTIA), BLUPs of severity from the leaf disk assay via vision (DSVI), BLUPs of severity from the leaf disk assay via image analysis (DSIA), mean of field evaluation of BLS in two locations (mean TL), highest score of BLS in NY (MNY), and highest score of BLS in two locations (MTL).

In 478-DTVI-DTIA-DSIA, the significant markers were on chromosomes 2K, 2N, 3K, 4N, 5K, 7K, 7N and 9N. These 14 SNP markers explained phenotypic variances for DTVI, DTIA and DSIA of 4.25, 1.77, and 22.62%, respectively (Table S5). In 4X-DSVI-MNY, there were three SNPs on chromosome 3K, 3N and 7N, accumulatively explaining 9.78% for DSVI and 11.31% for MNY. In 8X-DTVI-DSIA, only one significant marker showed on chromosome 9N explaining 0.64% of DTVI and 11.31% of MNY. In low-mean TL, the eight significant markers were on chromosomes 1K, 5N, 6N, 7N, and 9N explaining 45% of mean TL. In up-DTIA-MTL, the significant markers were on chromosome 1K and 7K explaining 5.03% of DTIA and 7.77% of MTL. The PVEs of each SNP in each subgroup and each trait are reported in Table S6.

In addition to the effect size of each SNP to each trait via PVE, the direction and inheritance of genotypic effects was shown in the format of allele combination effect of each SNP on phenotypes (Figure 8 and S10-S17). Expectedly, none of the SNPs showed inheritance with all of phenotypes, but with only some phenotypes. Most of inheritance patterns were dominant (Figure 8a, b, c, d, f, g, m, n, o, and q). A few were over dominant (Figure 8e, h, j, and k). Only Chr07N.48546809 showed and additive effects to DSIA as it contributes to the trait with PVE at 2.31 (Figure 8i). However, the PVE did not explain PVE. For example, Chr07N.61547213 had PVE 3.67 for MNY but did not show inheritance (Figure 8p).

**Figure 8.**
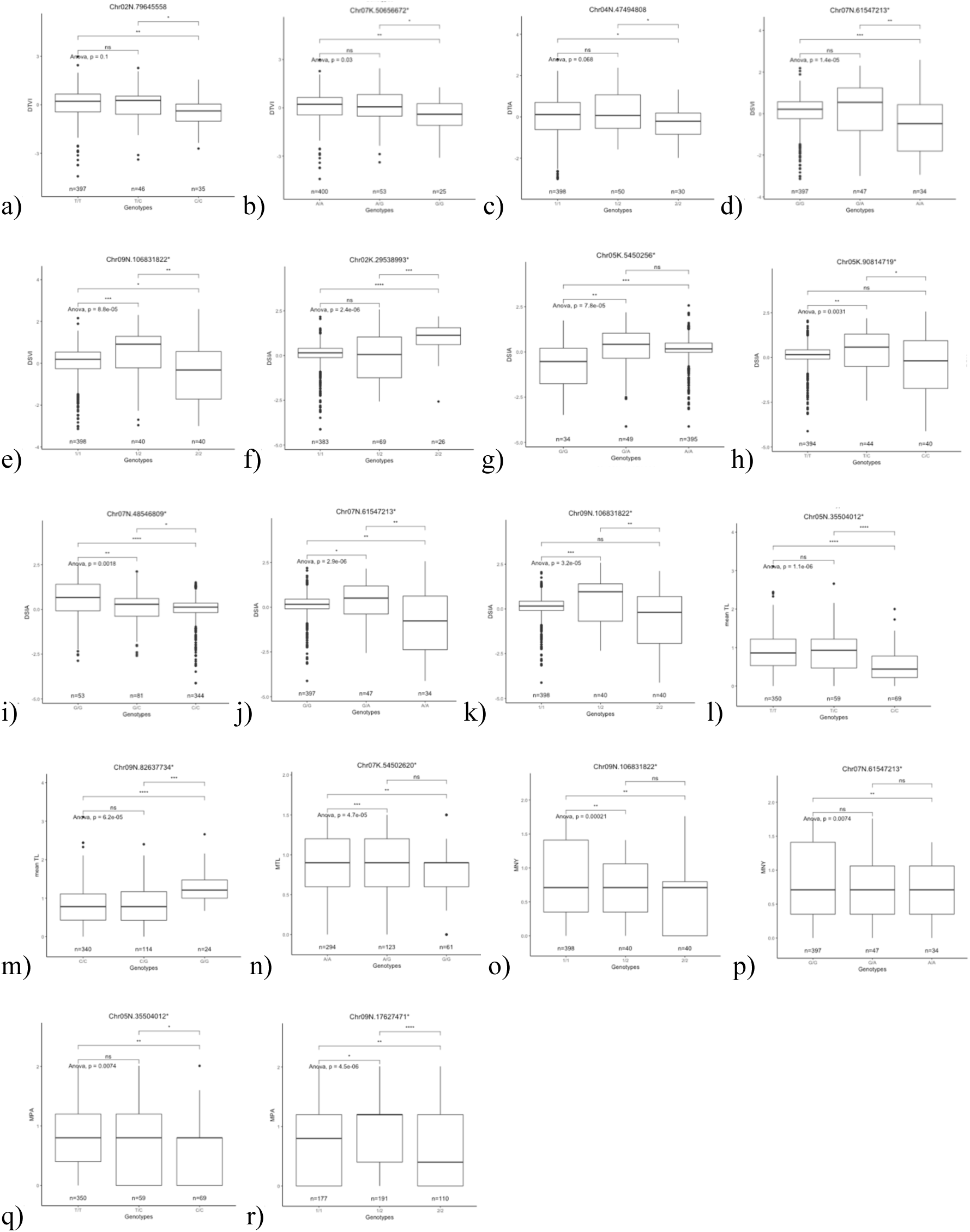
Examples of changes in DTVI, DTIA, DSVI, DSIA, mean TL, MTL, MNY and MPA associated with alleles at selected candidate single nucleotide polymorphisms (SNPs) in 478 genotypes. The effects of three different allele combinations were tested at alpha = 0.05. “ns” means non-significant, “*” means significant at alpha 0.05, “**” means significant at alpha 0.01, and “***” means significant at alpha 0.001.

When comparing these significant SNP markers to the potential candidate genes for resistance to BLS in rice, Chr07N_48546809, on chromosome 7N from 478-DTVI-DTIA-DSIA linking with Pavir.7NG240300 overlapped with LOC_Os04g39430.1 on rice chromosome 4 from AE11005627 marker (Table S1). Moreover, Chr07_54502620, on chromosome 7K from up-DTIA-MTL linking with Pavir.7KG238800 overlapped with LOC_Os04g43730 on rice chromosome 4 from ad04009558 marker.

## DISCUSSION

### The most resistant populations for improvement of resistance to BLS

Since the various phenotyping approaches yielded various resistant populations, it was challenging to determine the most resistant candidates for further breeding. Based on field evaluation, BLUPs from two locations were zero due to zero broad-sense heritability. The different trend for BLS resistance between two locations was confirmed by the significant rank test. This re-ranking incidence suggested that the resistance was under GxE effects between two locations (Le et al., 2001; Muir et al., 1992). If the mean of field scores across the two locations were used, the upland Shelter should be considered the most resistant population across two locations with a mean of 0.44. The high resistance of Shelter from the field evaluations could explain the result in a prior study (Songsomboon, 2019) that it cannot be improved for resistance to BLS via recurrent phenotypic selection in seedlings. However, the one of most susceptible SW787 had the mean score only 2.33. This suggested the low natural inoculation in 2017, which was not conducive to severe disease development. When comparing each location, SW803 showed lowest severity in NY but not significantly different than the other 55 genotypes, but it had highest score in PA at 2.11, which was significantly different. Therefore, this variation needs to be confirmed with more replicates, years, and locations. One more location in New Jersey was transplanted in 2018.

To lessen the GxE effect on the BLS, artificial inoculation was conducted in the laboratory. In the detached leaf assay, due to the limitation of sampling, a whole set of 478 genotypes cannot be tested at the same time. The experiment was separated into seven overlapping sets. However, there was a significant variation among sets. Such a variation can occur from the different age of each genotype in the same sampling time. Although the same stage of leaves was sampled at 30 centimeters from the top, variation existed among leaves from the same plant. To handle the variation among sets and unbalanced design (Hill & Rosenberger, 1985; Piepho et al., 2008; Robinson, 1991; Stroup & Mulitze 1991; White & Hodge, 2013), BLUPs were used by fitting the effect from the set as a fixed effect. Another artificial inoculation was leaf disk assay, with less limitation of the experimental size, the whole set of 478 genotypes was sampled at the same time with resistant and susceptible checks in each plate. However, this experiment was conducted only once at the end of growing season. Although a small effect of plates existed, BLUPs was fitted by considering plates as a fixed effect. Both artificial inoculations were evaluated via vision and image analysis. The image analysis is better regarding repeatability and reanalysis in the future (Stewart and McDonald, 2014). For the sake of comparison for the most resistant population, mean of percent lesion from image analysis was considered. In the detached leaf approach, the lowest severity population from DTIA was ECS.6 at 8% (not significantly different than the other 26 populations), but the population appeared very susceptible from DSIA at 94%. In the leaf disk assay, the most resistant population from DSIA was SW803 with 20 other populations. These supported the low correlation among all three disease evaluation approaches. The low correlation between laboratory and field evaluations could be explained by the different disease pressure between the two conditions. The natural inoculation in the field varied by the weather condition of each year whereas the laboratory inoculation was conducted with the high inoculum concentration to maximize the disease pressure. Another concern was the low correlation between detached leaf assay and leaf disk assay. It can be the result of the different ratio of leaf area to inoculum. The detached leaf assay used 5-cm long whole leaf and spraying inoculum whereas the leaf disk assay used only 8-mm bored leaf section with a 2-microliter inoculum droplet on a single location. The difference between spraying and droplet also showed in the assessment of early blight (*Alternaria solani*) (Chaerani et al., 2017).

The Qst-Fst of all phenotypes showed that none of them has gone under positive selection across five ecotypes. This neutrality was not expected as most of the resistance genes were hypothesized to be under natural selection (Grant et al. 1995, 1998; Stahl et al. 1999; Tian et al. 2002, 2003). However, neutral evolution of resistance genes like nucleotide binding sites – leucine rice repeats (NBS-LRR) can be undergone in a relaxed selective constraint. For example, when pathogens are absent or at low frequency, the resistance genes may experience much weaker or no cost of resistance (Gos and Wright, 2008). Such a low disease pressure can also be noticed by the means field evaluation that the highest severity was only 2.33 from 5 (the most severe).

### The improvement of multi-trait GWAS from single trait GWAS

Although all single traits from a different phenotype approach had moderate to high heritability, none of them provided significant peaks from GWAS from theoretically expected QQ-plots that were not distributed as expected from theory. This incidence can be explained by the high density of markers with high collinearity between markers and too small population size, giving the low power detection (Muir, 2007). Moreover, to utilize the SNP from multiple phenotype, multi-trait GWAS was suggested than aggregating the association SNP from each single phenotype (Cotsapas et al., 2011). Although the low correlations among phenotyping results of BLS resistance, such a low correlation or even negative correlation can lead to increase in effect size of SNP and thus increase statistical power, as happened with the tetraploid group by combining DSVI and MNY (Porter & O’Reilly, 2017; Zhu et al., 2015). On the other hand, combining highly positively correlated traits can reduce the average effect size and power as happened in the upland group combining DTVI and DTIA. However, the addition of traits did not always improve the analysis. In the lowland group, for example, only mean of the two locations yielded the most significant peaks of GWAS with theoretically expected QQ-plot.

Despite the improvement of multi-trait GWAS over single-trait GWAS, 28 SNP markers can explain the phenotypic variance ranging from only 3.28 to 26.52% among DTVI, DTIA, DSVI, DSIA, mean TL, MNY and MTL. The low PVE from GWAS is a common phenomenon and generally caused by missing heritability and can be improved by more genomic imputation and implementation of a large-scale next-generation sequence data (Fan and Song, 2016).

Moreover, each genotype group can show significant peaks from GWAS with specific traits. For example, the tetraploid group only provided the significant peaks from GWAS with DSVI-MNY whereas the full set of 478 genotypes provided the best fit GWAS with DTVI-DTIA-DSIA. To further study resistance to BLS in switchgrass populations, single-trait GWAS is not suggested due to the low power of determining significant markers (Porter & O’Reilly, 2017; Zhu et al., 2015). Multi-trait GWAS from both artificial and natural inoculation can help to improve the power.

### Dissecting resistance to BLS

From the QTLs study of Sato et al. (2015), the candidate resistance genes in rice were reported to be on three chromosomes with 13 SNPs markers within the LD of 200 kb (Table S1). There were 278 genes linked to these markers. Among them, 172 genes were annotated with 134 biological functions. There were 106 non-annotated genes. The most frequent biological functions were Leucine-rich repeat family protein (6), peroxidase precursor (6), transposon protein (6), nucleotide binding sites – leucine rice repeats (NBS-LRR) type disease resistance protein (5), MYB family transcription factor (3), dirigent (2), pentatricopeptide repeat (PPR) domain (2), fasciclin domain-containing protein (2), flavin monooxygenase (2), and heavy metal associated domain (2).

There were only two significant SNP markers that overlap with candidate resistance genes from rice Chr07N_48546809 from the 478 GWAS and Chr07K_54502620 from upland GWAS (Table 1 and S7). Chr07N_48546809 linked to Pavir.7NG240300 encoding for cytochromes P450, which involved diverse oxidation reactions and triterpene synthesis (Geisler et al.,). The most well-known triterpene against fungal disease was Avenacins in oat (*Avena* spp.). Chr07K_54502620 linked to Pavir.7KG238800 encoding for EGF_CA which has been proved to be the main structure of *ZmWAK* conferring resistance to maize head smut by controlling the galacturonan-binding for cell wall mediation (Zuo et al. 2015). In addition to genes overlapping with the resistance rice genes, Chr07N_61547213 showed as the significant peak in both 478 and 4X GWAS linked with Pavir.7NG322400 encoding for PPR repeat, which is one of the potential resistance genes in rice. These PPR repeat families were commonly known for disease resistance genes (Geddy & Brown, 2007). Pavir.2NG397900 encoding GDP-fucose protein O-fucosyltransferase relating to cell wall development and disease resistance (Perrin et al., 1999).

Although the rest of significant peaks are unique within each group of GWAS, we considered the biological functions for the resistance over the fixation of the alleles in each population for better resistance mechanism. In the full set of 478, on chromosome 2N, Pavir.2KG226200 was predicted to function as a zinc-binding RING finger. The zinc finger domains suggested the broad-spectrum nature of rice blast resistance gene *Pi54* classed as NBS-LRR, which played an important role in effector-triggered immunity (ETI) (Gupta et al., 2012). Moreover, this gene also encoded glycosyltransferase, which was responsible for cell wall synthesis and modification leading to disease resistance (Vorwerk et al., 2004). Pavir.2NG424700 encoded for pyruvate dehydrogenase, which is a key protein in tricarboxylic acid cycle. This can suggest the high energy intensive for resistance response as happened in wheat leaf rust (*Puccinia triticina* Eriks) (Bolton et al., 2008). Pavir.2NG500000 encoded for casein kinase II. The protein itself did not confer the resistance but it is often found as many of motifs in Mlo (Niu & He, 2009) which was identified to provide broad resistance to powdery mildew in barley (Jorgensen, 1992). On chromosome 3K, Pavir.3KG329700 encoded DNA polymerase III mainly known for DNA replications and repairing DNA damaged by hydrogen peroxide (Hagensee et al., 1987). The DNA repair process is part of the plant immune response (Song et al., 2011) especially in the infection of *B. oryzae* that hydrogen peroxide was shown to accumulate in a leaf (Kim et al., 2014). Pavir.3KG358400 encoded for serine aminopeptidase conferring no resistance but playing a major role in necrosis (López-Otín & Bond, 2008). On chromosome 4N, since Chr04N_47494808 was more closely linked to Pavir.4NG267200 encoding for uncharacterized protein suspected of hydrolase activity, the next Pavir.4NG267500 was considered for its function. The further gene encoded for Villin which is a tissue-specific actin modifying protein responsible for anti-apoptotic activity suppressing necrosis (Khurana & George, 2008; Xu et al., 2004). On chromosome 5N, Pavir.5KG028600 encoded for tetratricopeptide repeat which was one of five domains of SGT1 resistance gene in Arabidopsis (Azevedo et al., 2002). Pavir.5KG482000 encoded for ATP-binding cassette (ABC) transporter. There are many subgroups of ABC. In Arabidopsis, the mutant lacking the ABC transporter of penetration3/pleotropic drug resistance8 (PEN3/PDR8) conferred nonhost susceptibility to *Blumeria graminis*, suggesting that the ABC transporter was involved in exporting secreted toxins from the fungus and fungal suppressor from the host (Stein et al., 2006). Also, in wheat, the ABC suggested the exportation of mycotoxin conferring resistance to Fusarium head blight (Walter et al., 2015). On chromosome 7K, Pavir.7KG196600 encoded for interleukin-1 receptor-associated kinase, which was the pathogen recognition resulting in plant response (Kumagai et al., 2008). Pavir.7KG255000 encoded for oligo peptide transporter protein, which was also shown in rice QTL resistance to rice blast (*Manaporthe grisea*) (Wang et al., 2001). Lastly for full set, on chromosome 9N, Pavir.9NG844200 encoded for Mitochondrial Fe(II) transporter MMT1. Despite no direct report of resistance from this activity, mitochondria were proven to control redox homeostasis under stress (Dutilleul et al., 2003).

In GWAS from 4X-DSVI-MNY, on chromosome 3K, Pavir.3KG095200 encoded for both peroxisomal membrane protein and ABC transporter. As the high involvement of ROS in the infection of *B. oryzae* (Antonious and Jawhar, 2013), peroxidase played an important role in Pathogen-associated molecular pattern-triggered immunity (PTI) (Mammarella et al., 2015). Additionally, Pavir.3NG101800 on chromosome 3N encoded ABC transporter.

The GWAS from 8X-DTVI-DSIA yielded only one significant SNP on chromosome 9N. Pavir.9NG061100 encoded for DNA topoisomerase and more importantly dirigent. Dirigent protein conferred resistance by lignin and lignan synthesis (Ralph et al., 2006). The lignin synthesis was expected to relate with resistance to BLS by strengthening cell walls (Dallagnol et al., 2013).

In GWAS from lowland mean of two locations, on chromosome 1K, the further linked Pavir.1KG225100 encoded for MORC family-ATPases. This protein is the key component of compromised for recognition of Turnip Crinkle Virus (CRT1) in Arabidopsis that are required for PTI, basal resistance, no-host resistance and systemic acquired resistance (Kang et al., 2012). On chromosome 5N, Pavir.5NG192000 encoded for COP9 signalosome complex. Interestingly, this signalosome complex interacts with SGT1, which Pavir.5KG028600 from GWAS of 478 genotype set encodes, indicating the disease resistance (Azevedo et al., 2002). Pavir.5NG248900 encoded basic leucine zipper (bZIP) transcription factor, which is a key modulator in hypersensitive response and salicylic acid (Alve et al., 2006). TB2/DP1, Pavir.5NG476900 encoded for abcisic-acid induced TB2/DP1, HVA22 family responsible for membrane turnover or decrease unnecessary secretion against Southern corn leaf blight by *Cochliobolus heterostrophus* (Drechs.) Drechs. in maize (Balint-Kurti et al., 2010). On chromosome 6N, Pavir.6NG083100 encoded homeobox-leucine zipper protein playing an important role in responses to abscisic acid under stress (Gago et al., 2002), transcriptional regulation for disease resistance in rice (Luo et al., 2005) and programmed cell death (Mayda et al., 1999). On chromosome 7N, Pavir.7NG091000 encoded sucrose synthase. Herbers et al., (1996) suggested that hexose sensing was important to activate defense-related genes as well as repression of photosynthetic genes. On chromosome 9N, Pavir.9NG173600 encoded Phytochrome-interacting factor 4. Photo receptors in plants plays essential role in hydrogen peroxide and ROS in the cell resulting in disease response (Karpinski et al., 2003). Lastly, Pavir.9NG499500 encoded for alpha/beta hydrolase which is a diversely biochemical protein family involving the activation of hydrogen peroxide (Lenfant et al., 2012).

In the last GWAS from upland DTIA-MTL, Chr01K_61692577 linked to two interesting genes. First, Pavir.1KG372200 encoded Exo70 exocyst complex subunit, which was required for recognizing the avirulent effector AVR-Pii from rice blast resulting in ETI (Fujisaki et al., 2015). The other linked gene was Pavir.1KG372200 encoding for Nicotinamidase that showed resistance to tomato mosaic virus (Arens et al., 2010).

Although these candidate genes from the switchgrass association studies shared some functions from candidate genes from rice resistance to BLS, some of the potential functions conferring resistance were still missing such as MYB. In rice, the *MYB* gene controls Jasmonic acid (JA) against rice blast fungus: *Pyricularia greisea* (Ambawat et al., 2013; Lee et al., 2001). In Arabidopsis interaction with necrotrophic *Alternaria brassicicola*, the response to JA was the upregulated plant defensin protein (*Pdf*) gene.

## CONCLUSIONS AND POTENTIAL APPLICATIONS

In conclusion, the resistance to BLS resulted differently based on different phenotyping approaches, with low correlations suggesting that resistance was sensitive to phenotyping methods in different conditions. Although it is difficult to standardize the phenotyping approach that can take account of all conditions, finer and more powerful phenotyping of the resistance is still needed. As such, to the best of our phenotyping combination, ‘SW788’, ‘SW806’, ‘SW802’, ‘SW793’, ‘SW781’, ‘SW797’, ‘SW798’, ‘SW803’, ‘SW795’, ‘SW805 should be considered as candidates for resistance. To increase the statistical power in SNP detection, the five subgroups of populations were conducted with multi-trait GWAS. The full population of 478 genotypes showed the most SNP markers that can be considered as potential candidate genes related to resistance to BLS with reasonable biological functions. Additionally, the SNP markers from the other subgroups provided broader resistance mechanism. Nevertheless, one needs to be prudent regarding the utilization of these markers. The dependency of multitrait GWAS to yield significant SNP peaks brought the challenge for genomic-assisted breeding for resistance to BLS. The application of these markers cannot be directly transferred for genomics-assisted selection without validation to the choices of phenotyping approaches in new populations.

When considering biological functions from the GWAS regardless of subgroups and trait combinations, the defense mechanisms against BLS are complicated and multifaceted, probably controlled by multiple small-effect alleles. The defense mechanism started from basal defense via cell-wall mediated proteins and PTI via peroxidase. Then, ETI depending on NBS-LRR, and PPR repeat took place. The responses to necrosis were suppressed by homeostasis and anti-apoptotic proteins. To our knowledge, this was first research to dissect the resistance to BLS in switchgrass. With the current power of detection within the population, multi-trait GWAS can provide functional insight into the resistance within multiple subgroups of genotypes.

## Supporting information

Post-hoc Tukey's HSD comparisons

Supplemental explanation

## ACKNOWLEDGEMENTS

The project is supported by PhD funding from the Development and Promotion of Science and Technology Talents Project (DPST), the Royal Thai Government. The northern switchgrass association panel and its HAPMAP data were provided by Dr. Michael Casler, Dr. Paul Grabowski and Dr. Guillaume Ramstein from US Dairy Farage Research Center, USDA-ARS. The genome-wide association analysis with multiple traits was consulted by Dr. Deniz Akdemir. Visual and digital disease evaluations were guided by Dr. Shawn Kenaley and Dr. Ethan Stewart.

## Notes

#### Summary of Updates

reanalyze with updated reference genome v4.1 all statistical comparisons update H2 table on different groups added admixture analysis for population structure added Qst-Fst added allele inheritance

