## Supplemental figures for "Genome-Wide Associations with Resistance to Bipolaris Leaf Spot (*Bipolaris oryzae* (Breda de Haan) Shoemaker) in a Northern Switchgrass Population (*Panicum virgatum* L.)"

1

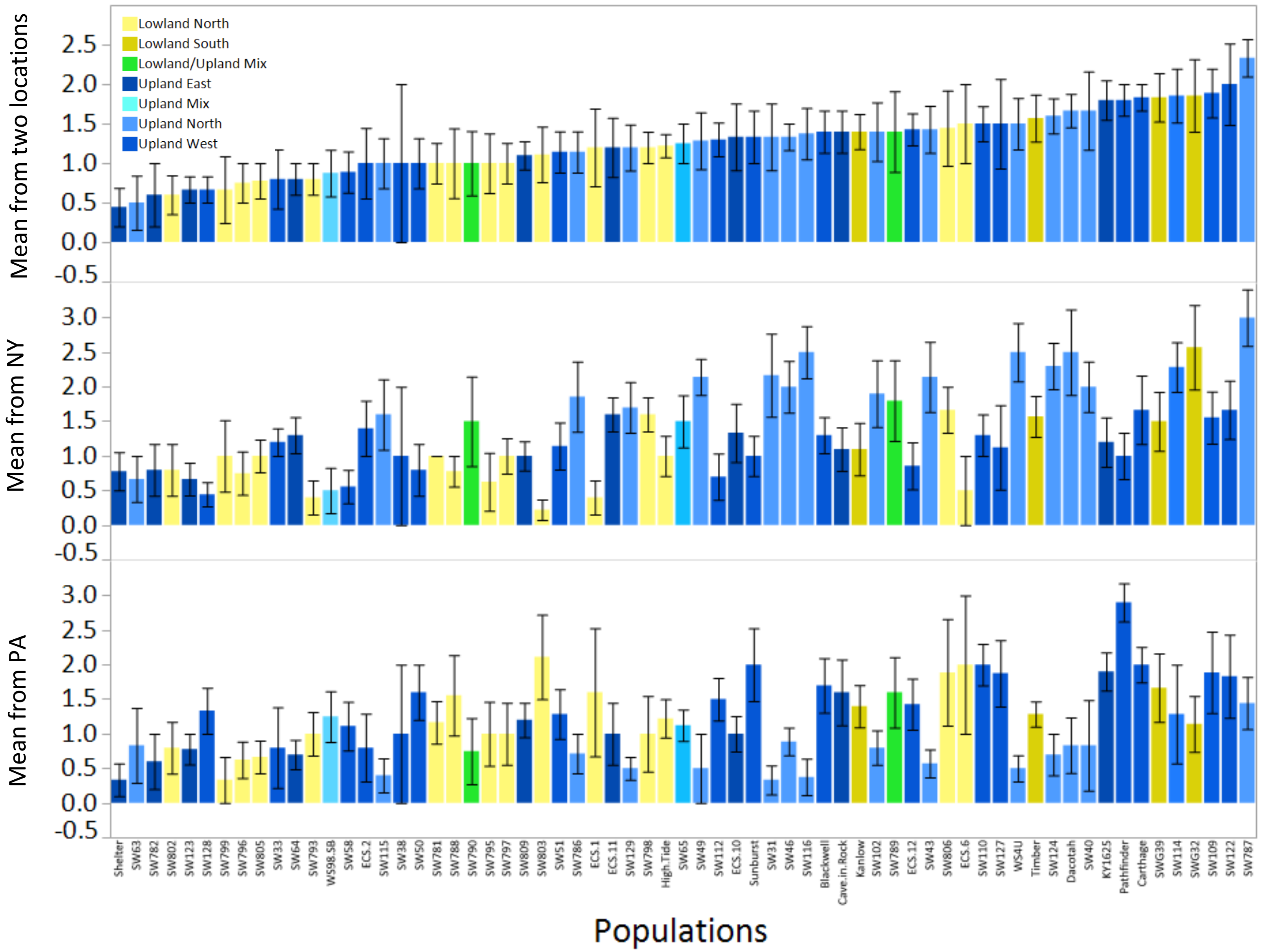


Figure S1. Histogram of mean severity score from 0 to 5 from two locations (top), mean from NY (middle) and mean from PA with standard error from each of the 66 populations. They were ordered based on the mean from the two locations and colored based on ecotypes.

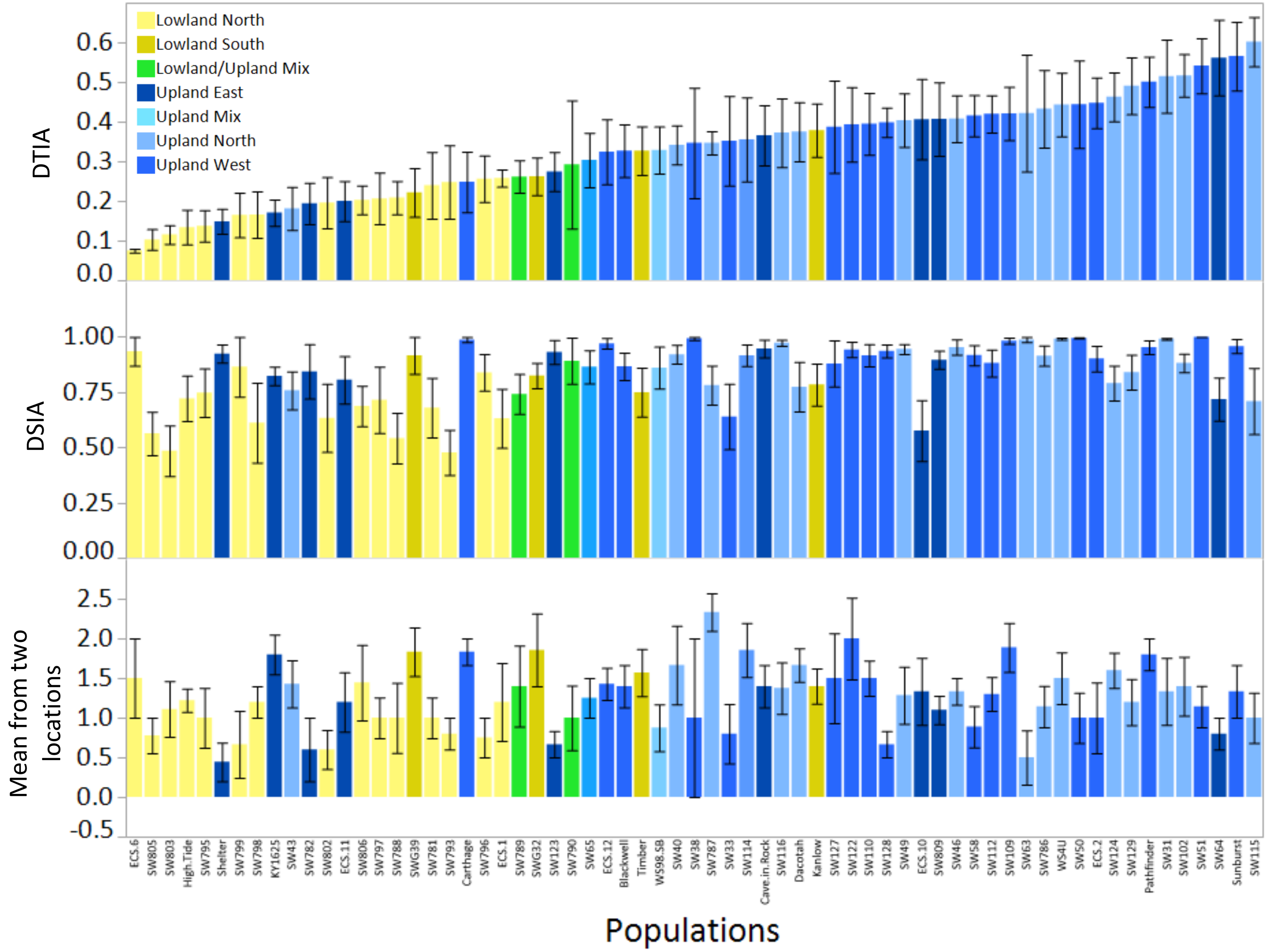


Figure S2 Histogram of mean severity from the leaf detachment assay via image analysis (DTIA) (top), from the leaf disk assay via image analysis (DSIA) (middle) and mean field severity from two locations with standard error from each of the 66 populations. They were ordered based on DTIA and colored based on ecotypes.

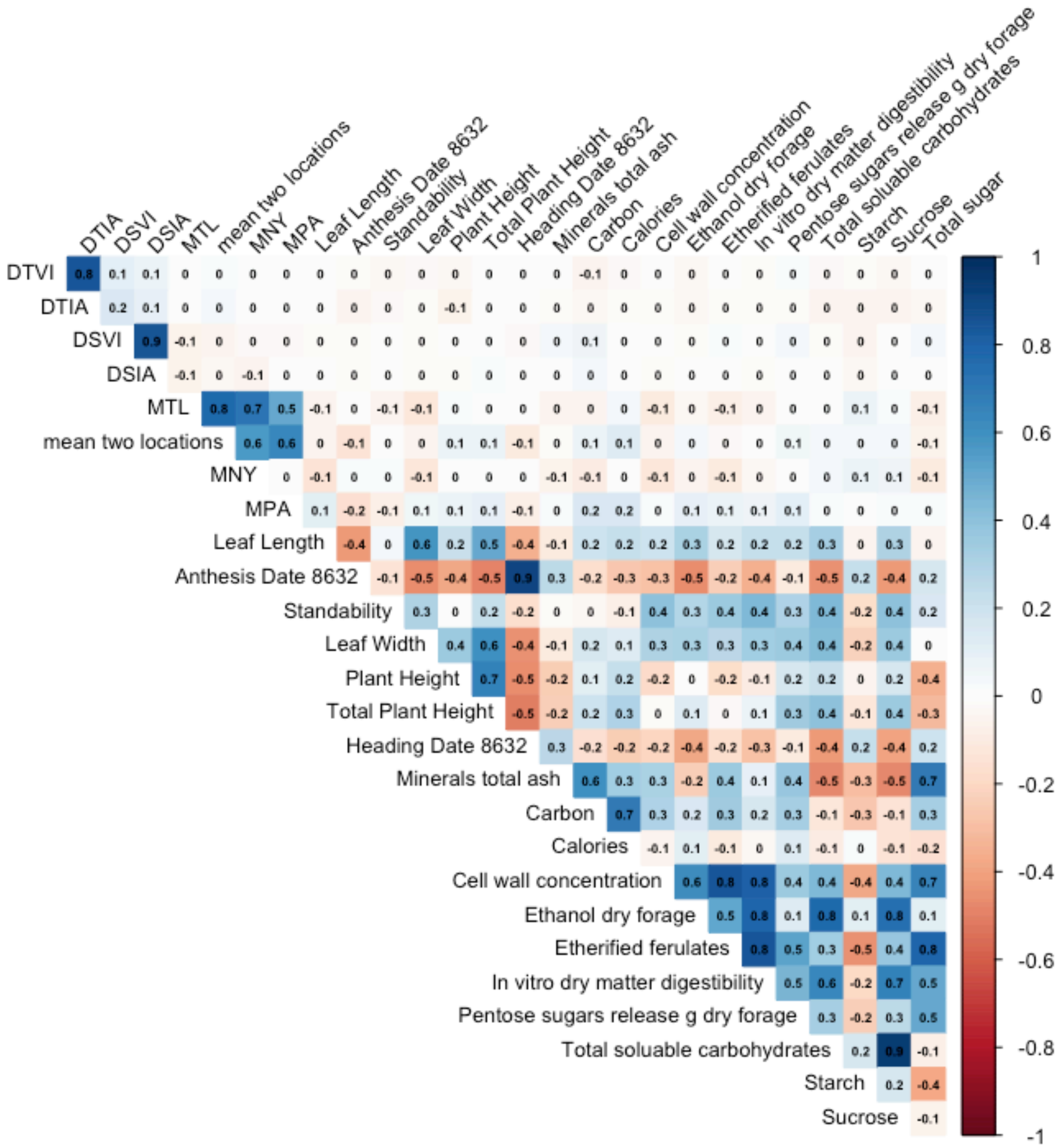


Figure S3. Correlation plot among BLUPs of severity from the leaf detachment via vision (DTVI), via image analysis (DTIA), from leaf disk assay via vision (DSVI), via image analysis (DSIA), the highest score between two locations (MTL), mean from two locations (mean TL), highest score in NY (MNY), highest score in PA (MPA) and BLUPs of other agronomic and biomass quality traits.

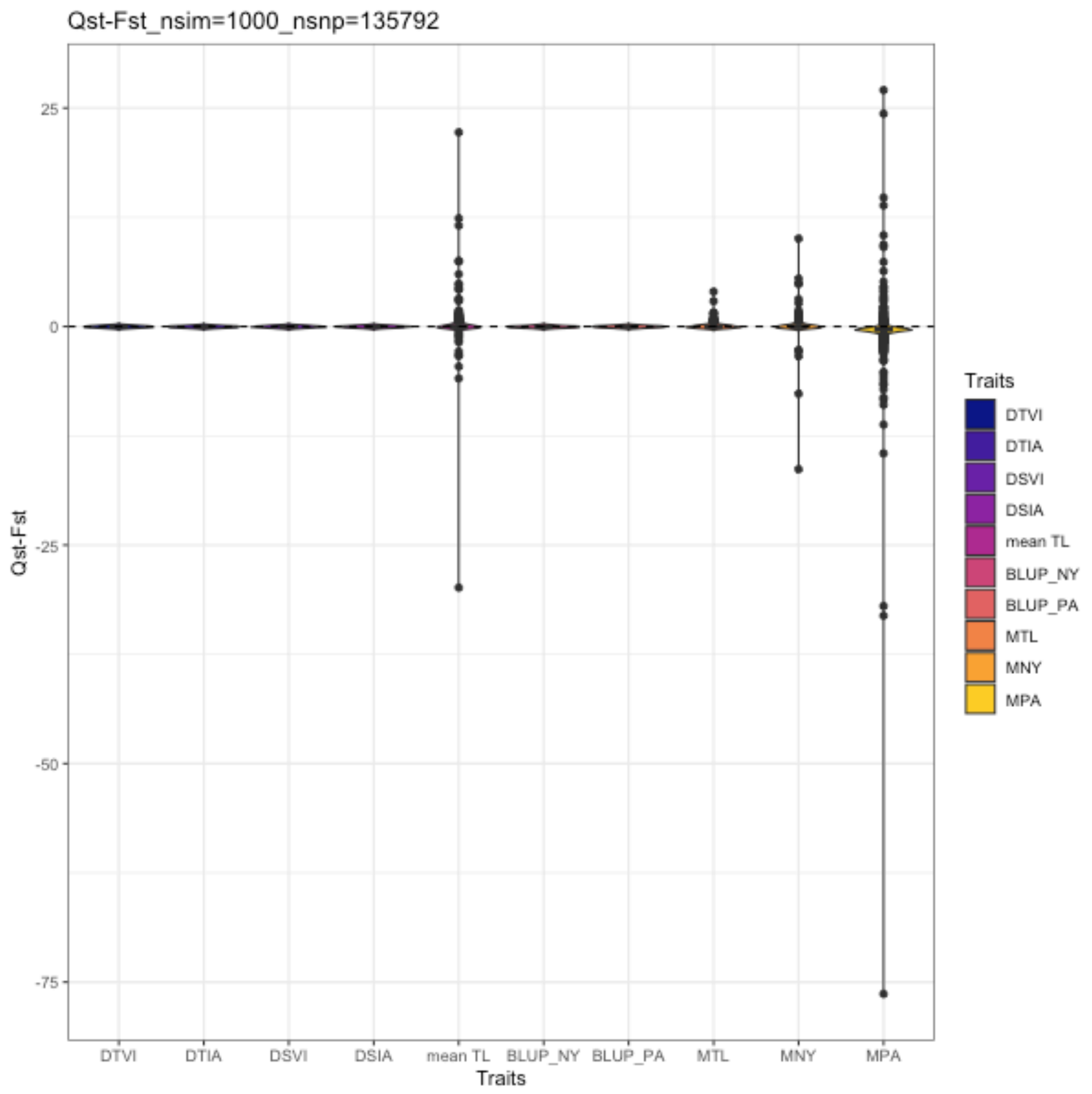


Figure S4. The bootstrapped distributions of Qst-Fst for each phenotypes (DTVI, DTIA, DSVI, DSIA, mean TL, BLUPs NY, BLUP PA, MTL, MNY, and MPA) is compared against the expected value of zero under neutrality (black dashed line).

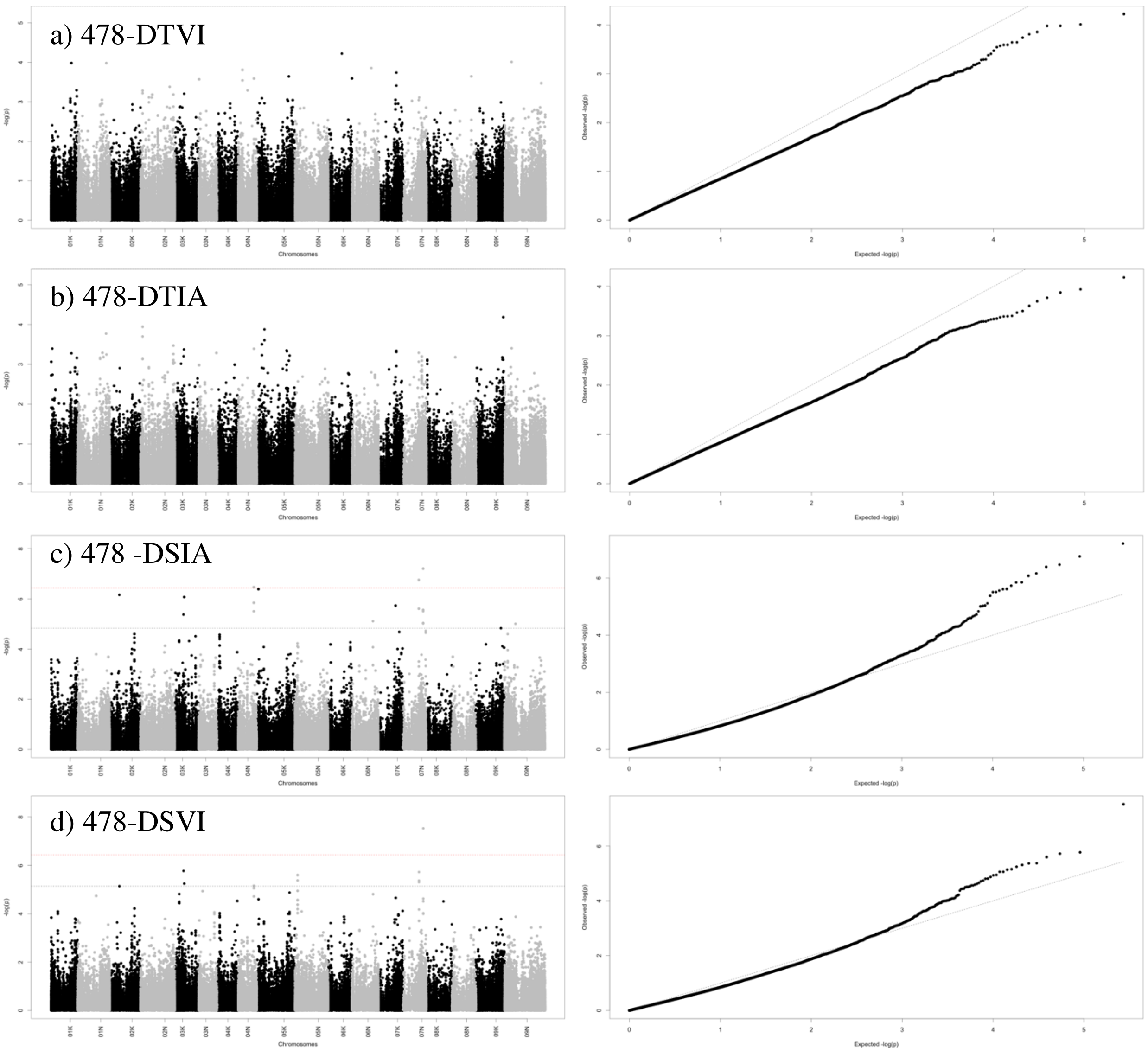


Figure S5. (Left) Manhattan plot showing genetic associated with a) DTVI, b) DTIA, c) DSIA, and d) DSVI in 478 genotypes. The black dashed line represents the FDR threshold (0.1) and the red dashed line represents Bonferroni correction threshold. On x-axis, the physical positions of the SNPs were aligned in 18 chromosomes of *P. virgatum*. (Right) Quantile-quantile (QQ) plots between the distributions of observed to expected P-values for GWAS of each trait combination in each genotype group.

Abbreviation: BLUPs of severity from the detached leaf via vision (DTVI), BLUPs of severity from the detached leaf via image analysis (DTIA), BLUPs of severity from the leaf disk via vision (DSVI), and BLUPs of severity from the leaf disk via image analysis (DSIA).

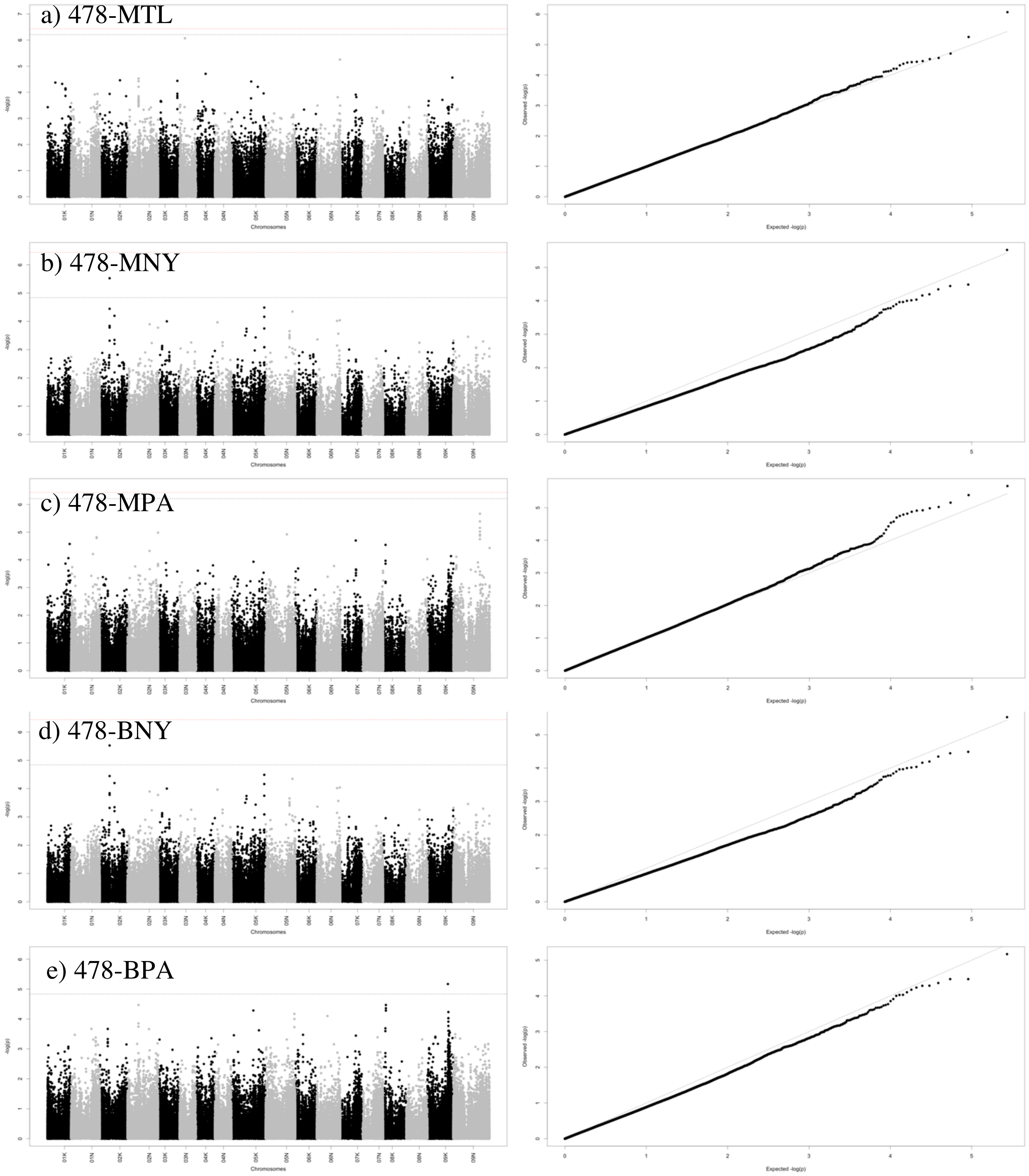


Figure S6. (Left) Manhattan plot showing genetic associated with a) MTL, b) MNY, c) MPA, d) BNY and e) BPA in 478 genotypes. The black dashed line represents the FDR threshold (0.1) and the red dashed line represents Bonferroni correction threshold. On x-axis, the physical positions of the SNPs were aligned in 18 chromosomes of *P. virgatum*. (Right) Quantile-quantile (QQ) plots between the distributions of observed to expected P-values for GWAS of each trait combination in each genotype group.

Abbreviation: Highest score of BLS in two locations (MTL), highest score of BLS in NY (MNY), highest score of BLS in PA (MPA), BLUPs of field evaluation in NY (BNY), and BLUPs of field evaluation in PA (BPA).

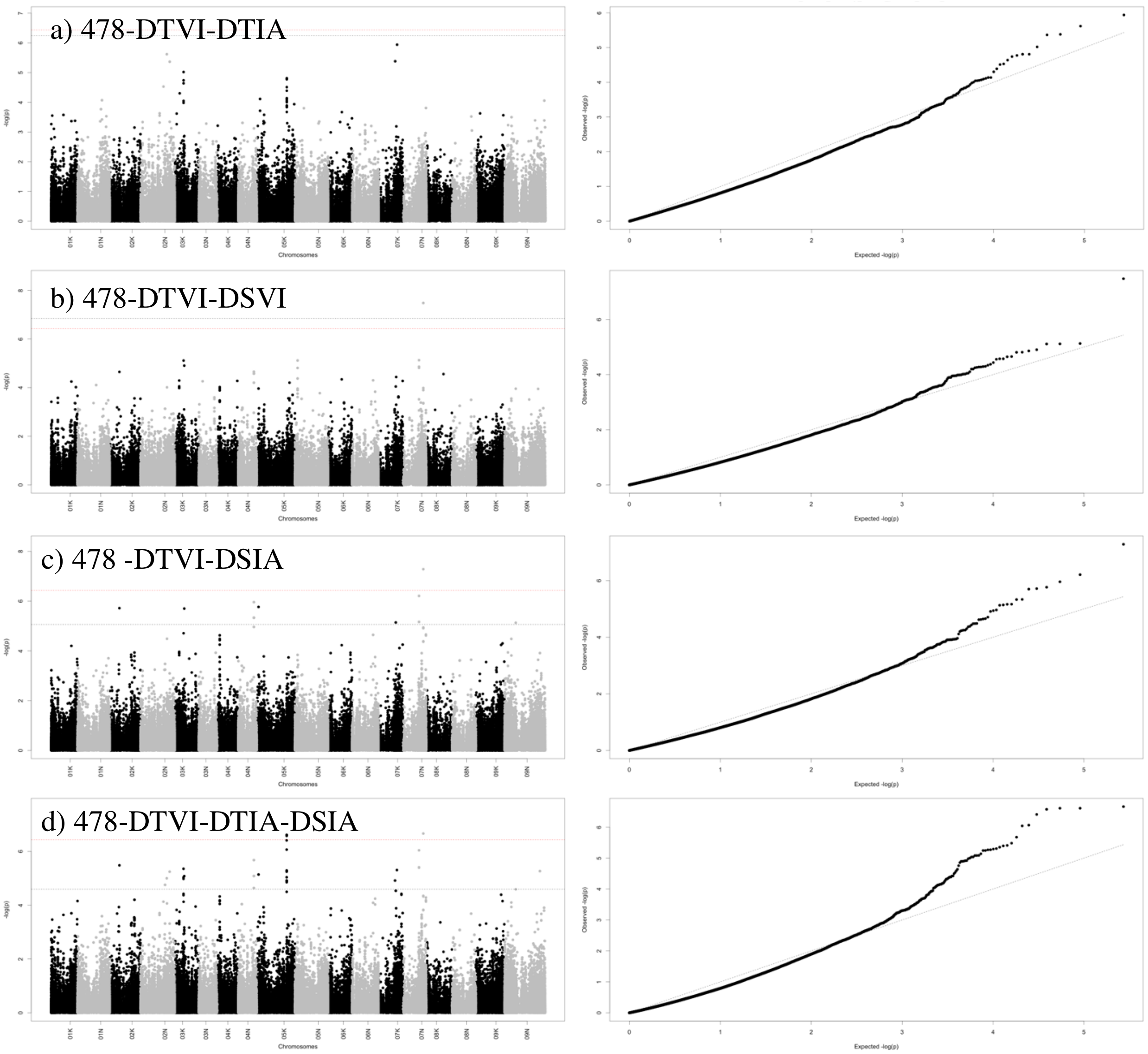


Figure S7. (Left) Manhattan plot showing genetic associated with a) DTVI, b) DTIA, c) DSIA and d) DTVI-DTIA-DSIA in 478 genotypes. The black dashed line represents the FDR threshold (0.1) and the red dashed line represents Bonferroni correction threshold. On x-axis, the physical positions of the SNPs were aligned in 18 chromosomes of *P. virgatum*. (Right) Quantile-quantile (QQ) plots between the distributions of observed to expected P-values for GWAS of each trait combination in each genotype group.

Abbreviation: Abbreviation: BLUPs of severity from the detached leaf via vision (DTVI), BLUPs of severity from the detached leaf via image analysis (DTIA), and BLUPs of severity from the detached leaf via image analysis (DTIA).

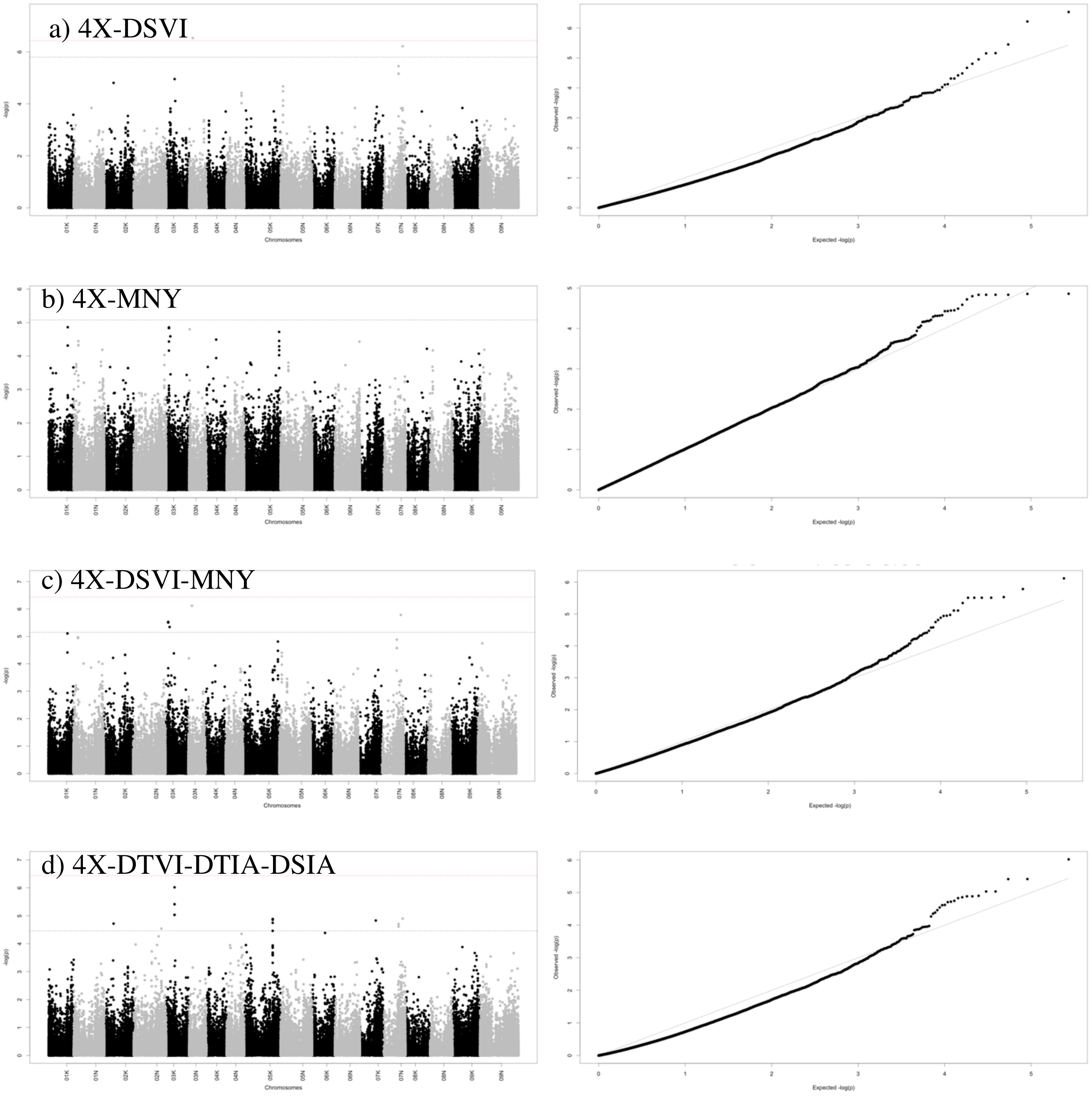


Figure S8. (Left) Manhattan plot showing genetic associated with a) DSVI, b) MNY, c) DSVI-MNY and d) DSVI-DTIA-DSIA in 4X genotypes. The dashed line represents the FDR threshold (0.1) and the line represents Bonferroni correction threshold. The black dashed line represents the FDR threshold (0.1) and the red dashed line represents Bonferroni correction threshold. On x-axis, the physical positions of the SNPs were aligned in 18 chromosomes of *P. virgatum*. (Right) Quantile-quantile (QQ) plots between the distributions of observed to expected P-values for GWAS of each trait combination in each genotype group.

Abbreviation: Abbreviation: BLUPs of severity from the detached leaf via vision (DTVI), BLUPs of severity from the detached leaf via image analysis (DTIA), BLUPs of severity from the leaf disk assay via vision (DSVI), BLUPs of severity from the leaf disk assay via image analysis (DSIA), and highest score of BLS in NY (MNY).

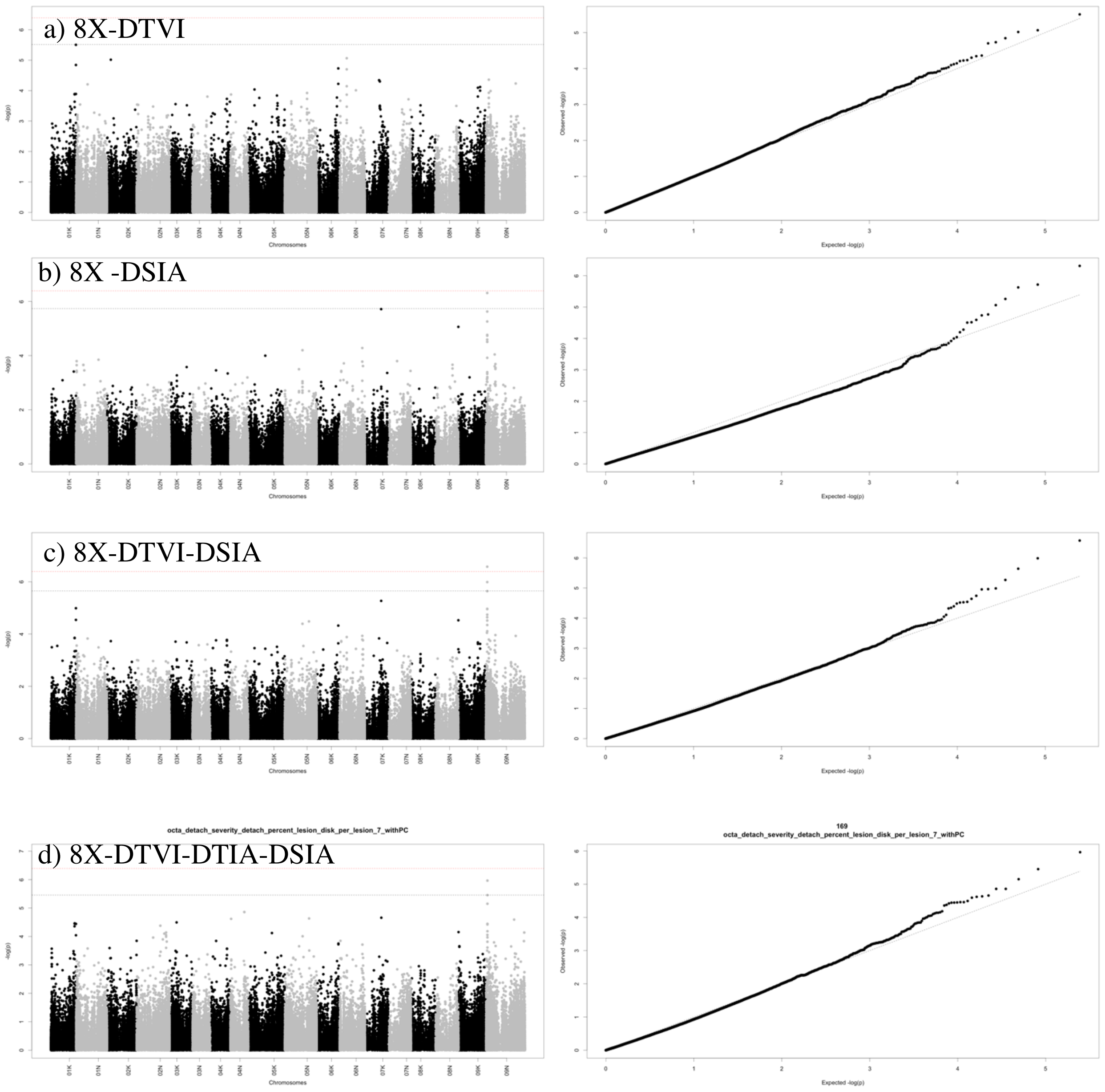


Figure S9. (Left) Manhattan plot showing genetic associated with a) DTVI, b) DSIA, c) DTVI-DSIA and d) DTVI-DTIA-DSIA in 8X genotypes. The black dashed line represents the FDR threshold (0.1) and the red dashed line represents Bonferroni correction threshold. On x-axis, the physical positions of the SNPs were aligned in 18 chromosomes of *P. virgatum*. (Right) Quantile-quantile (QQ) plots between the distributions of observed to expected P-values for GWAS of each trait combination in each genotype group.

Abbreviation: Abbreviation: BLUPs of severity from the detached leaf via vision (DTVI), BLUPs of severity from the detached leaf via image analysis (DTIA), and BLUPs of severity from the leaf disk assay via image analysis (DSIA).

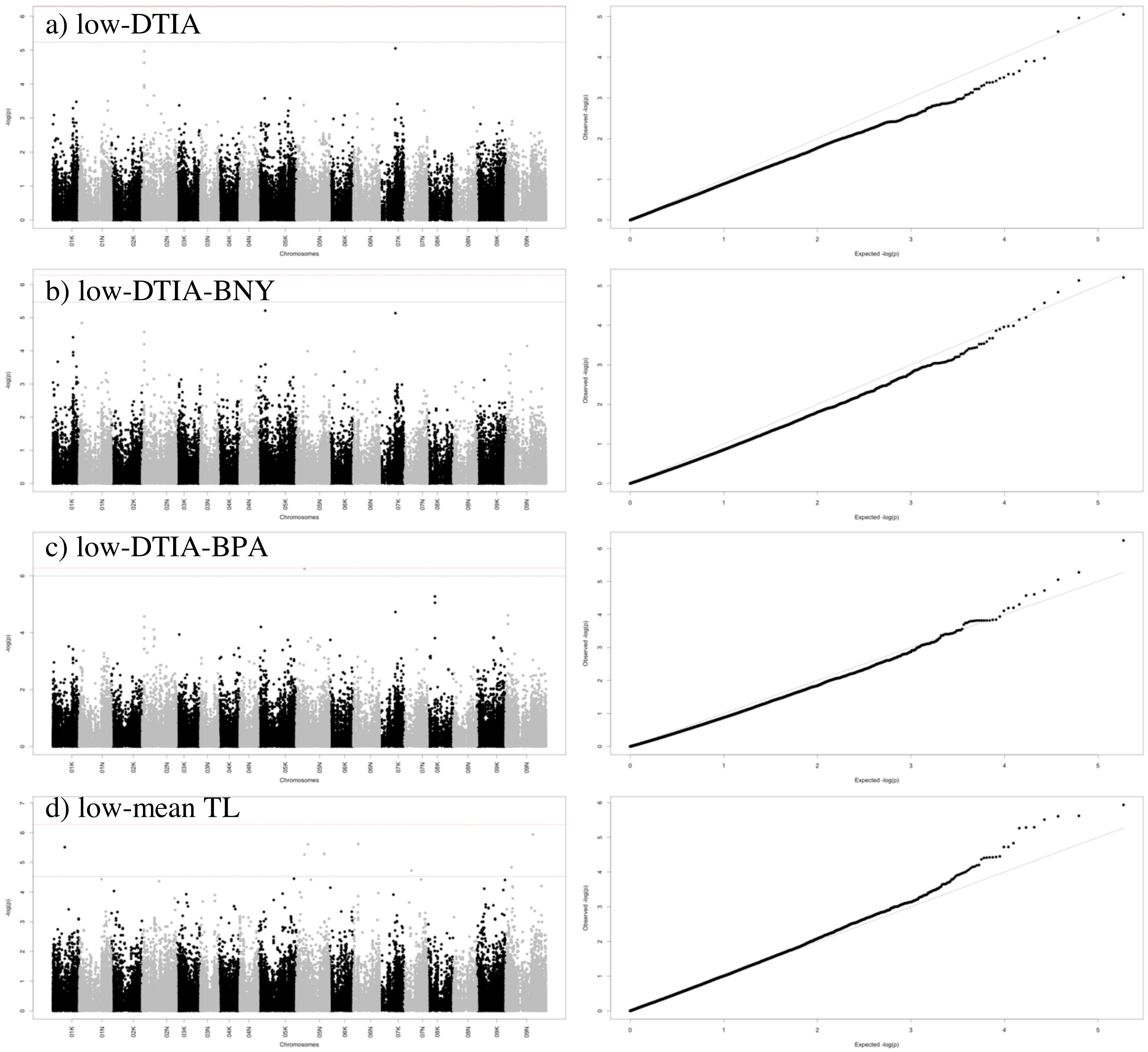


Figure S10. (Left) Manhattan plot showing genetic associated with a) DTIA, b) DTIA-NY, c) DTIA-PA and d) mean TL in lowland genotypes. The black dashed line represents the FDR threshold (0.1) and the red dashed line represents Bonferroni correction threshold. On x-axis, the physical positions of the SNPs were aligned in 18 chromosomes of *P. virgatum*. (Right) Quantile-quantile (QQ) plots between the distributions of observed to expected P-values for GWAS of each trait combination in each genotype group.

Abbreviation: Abbreviation: BLUPs of severity from the detached leaf via image analysis (DTIA), BLUPs of field evaluation in NY (BNY), BLUPs of field evaluation in PA (BPA), and mean of field evaluation of BLS in two locations (mean TL).

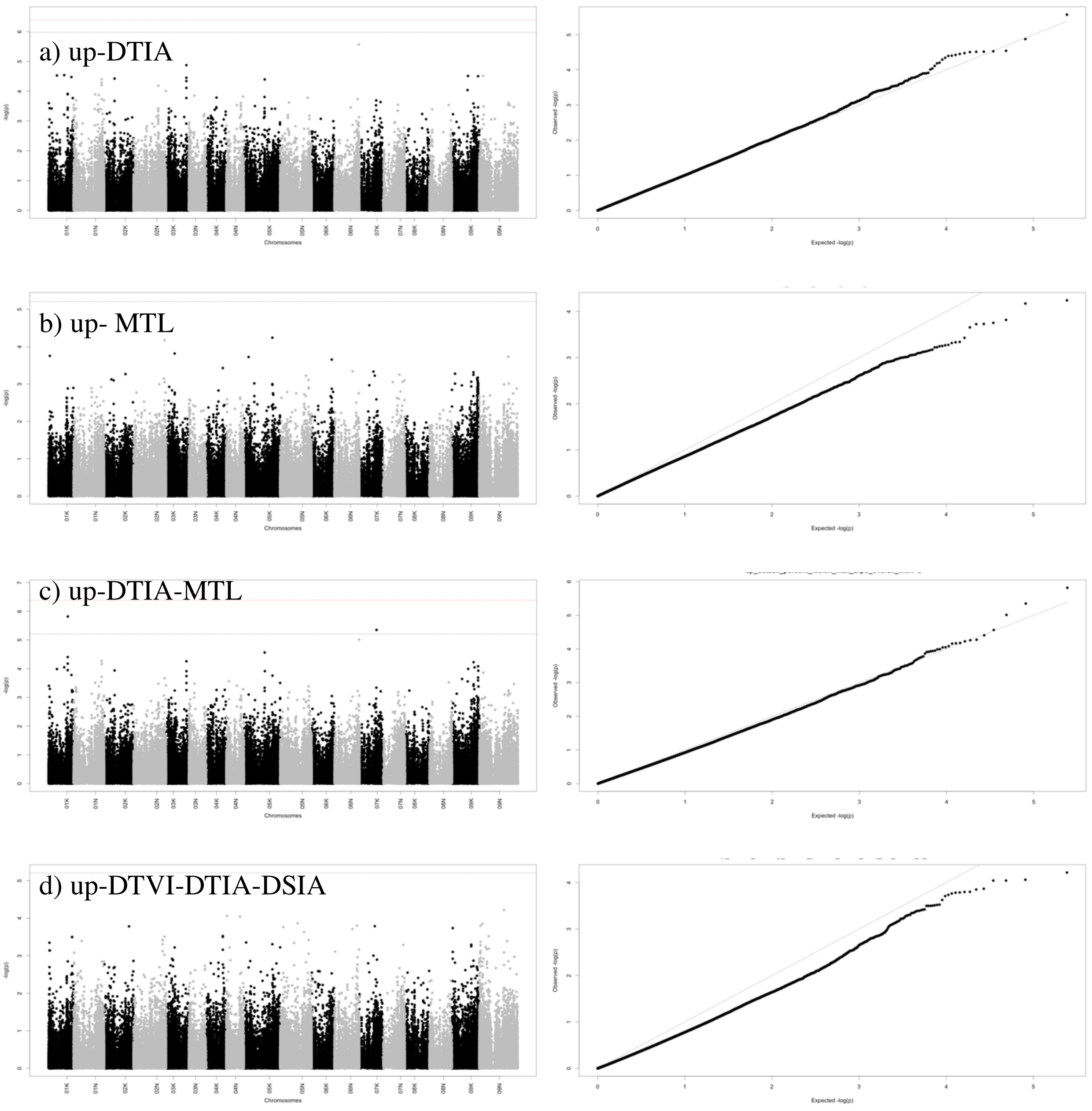


Figure S11. Manhattan plot showing genetic associated with a) DTIA, b) MTL, c) DTIA-MTL and d) DTVI-DTIA-DSIA in upland genotypes. The black dashed line represents the FDR threshold (0.1) and the red dashed line represents Bonferroni correction threshold. On x-axis, the physical positions of the SNPs were aligned in 18 chromosomes of *P. virgatum*. (Right) Quantile-quantile (QQ) plots between the distributions of observed to expected P-values for GWAS of each trait combination in each genotype group.

Abbreviation: Abbreviation: BLUPs of severity from the detached leaf via vision (DTVI), BLUPs of severity from the detached leaf via image analysis (DTIA), BLUPs of severity from the leaf disk assay via image analysis (DSIA), and highest score of BLS in two locations (MTL).
