## Supplemental explanation for "Genome-Wide Associations with Resistance to Bipolaris Leaf Spot (*Bipolaris oryzae* (Breda de Haan) Shoemaker) in a Northern Switchgrass Population (*Panicum virgatum* L.)"

**Java code for ImageJ analysis for detached leaf and leaf disk assay**

**For each leaf**

```
dir= getDirectory("");
// Get files in directory:
	files= getFileList(dir);
	path = dir + files[k];
	open(path);
	run("Rotate 90 Degrees Right");
//decode QR
	makeRectangle(414, 216, 990, 996);
	run("Duplicate...", " ");
	run("8-bit");
	//run("Brightness/Contrast...");
	setMinAndMax(204, 206);
	run("Apply LUT");
	run("QR Decoder", "error=FAILED");
	selectWindow("QR Code");
	QR_ID = getInfo("window.contents");
	run("Close");
	print(QR_ID);
	run("Close");
//analyze lesions on the leaf
	selectWindow(files[k]);
	makeRectangle(1476, 336, 5010, 1032);
	run("Duplicate...", " ");
	run("Duplicate...", " ");
	run("8-bit");
	setAutoThreshold("Default");
	//run("Threshold...");
	setThreshold(0, 188);
	//setThreshold(0, 188);
	setOption("BlackBackground", false);
	run("Convert to Mask");
	run("Analyze Particles...", "size=0-Infinity display include summarize");
	close();
	selectWindow("Summary");
	lines = split(getInfo(), "\n");
	headings = split(lines[0], "\t");
	values = split(lines[lengthOf(lines)-1], "\t");
	 for (i=0; i<headings.length; i++)
	     print(headings[i]+": "+values[i]);
	// Color Thresholder 1.51j
	// Autogenerated macro, single images only!
	min=newArray(3);
```

```
max=newArray(3);
filter=newArray(3);
a=getTitle();
run("HSB Stack");
run("Convert Stack to Images");
selectWindow("Hue");
rename("0");
selectWindow("Saturation");
rename("1");
selectWindow("Brightness");
rename("2");
min[0]=0;
max[0]=35;
filter[0]="pass";
min[1]=0;
max[1]=255;
filter[1]="pass";
min[2]=0;
max[2]=210;
filter[2]="pass";
filter[0]="pass";
filter[1]="pass";
filter[2]="pass";
for (i=0;i<3;i++){
  selectWindow(""+i);
  setThreshold(min[i], max[i]);
  run("Convert to Mask");
  if (filter[i]=="stop")  run("Invert");
}
imageCalculator("AND create", "0","1");
imageCalculator("AND create", "Result of 0","2");
for (i=0;i<3;i++){
  selectWindow(""+i);
  close();
}
selectWindow("Result of 0");
close();
selectWindow("Result of Result of 0");
rename(a);
// Colour Thresholding-------------

run("Analyze Particles...", "display include summarize");
close();

selectWindow("Summary");
lines = split(getInfo(), "\n");
```

```
headings = split(lines[0], "\t");
values = split(lines[lengthOf(lines)-1], "\t");
 for (i=0; i<headings.length; i++)
      print(headings[i]+": "+values[i]);
```

**Code for leaf disk assay**

```
//extract each disk from 28 disk in each plate
//run("Subtract Background...", "rolling=60");
// Color Thresholder 1.51j
// Autogenerated macro, single images only!
min=newArray(3);
max=newArray(3);
filter=newArray(3);
a=getTitle();
run("HSB Stack");

run("Convert Stack to Images");
selectWindow("Hue");
rename("0");
selectWindow("Saturation");
rename("1");
selectWindow("Brightness");
rename("2");
min[0]=0;
max[0]=126;
filter[0]="pass";
min[1]=150;
max[1]=255;
filter[1]="pass";
min[2]=56;
max[2]=255;
filter[2]="pass";
filter[0]="pass";
filter[1]="pass";
filter[2]="pass";
for (i=0;i<3;i++){
  selectWindow(""+i);
  setThreshold(min[i], max[i]);
  run("Convert to Mask");
  if (filter[i]=="stop")  run("Invert");
}
imageCalculator("AND create", "0","1");
imageCalculator("AND create", "Result of 0","2");
for (i=0;i<3;i++){
  selectWindow(""+i);
```

```
          close();
        }
        selectWindow("Result of 0");
        close();
        selectWindow("Result of Result of 0");
        rename(a);
        // Colour Thresholding------------
        run("Analyze Particles...", "size=1000-Infinity circularity=0.00-1.00 include clear include
        add");
        //try to rename
        //for ( i =0; i < roiManager("count"); i++) {
            //roiManager ("select", i);
            //yyyy=  split( call( "ij.plugin.frame.RoiManager.getName", i ), "-" );
            //roiManager( "Rename",yyyy[1] );
        //}
        //roiManager("Sort");
        close();
        for (i=0; i<roiManager("count"); ++i) {
        //Select the original which should be cropped
           run("Duplicate...", "title=crop");
           roiManager("Select", i);
           run("Crop");
           saveAs("[file_name]".tif");
         }
//analyze each disk
        ////////////////////////////////////////////total area
        showProgress(k, files.length);
              IJ.redirectErrorMessages();
              open(path);
              if (nImages>=1) {
        //remove background
        setBackgroundColor(0, 0, 0);
        run("Clear Outside");


        // Color Thresholder 1.51j
        // Autogenerated macro, single images only!
        min=newArray(3);
        max=newArray(3);
        filter=newArray(3);
        a=getTitle();
        run("HSB Stack");
        run("Convert Stack to Images");
        selectWindow("Hue");
        rename("0");
        selectWindow("Saturation");
```

```
rename("1");
selectWindow("Brightness");
rename("2");
min[0]=0;
max[0]=255;
filter[0]="pass";
min[1]=0;
max[1]=255;
filter[1]="pass";
min[2]=1;
max[2]=255;
filter[2]="pass";
filter[0]="pass";
filter[1]="pass";
filter[2]="pass";
for (i=0;i<3;i++){
  selectWindow(""+i);
  setThreshold(min[i], max[i]);
  run("Convert to Mask");
  if (filter[i]=="stop")  run("Invert");
}
imageCalculator("AND create", "0","1");
imageCalculator("AND create", "Result of 0","2");
for (i=0;i<3;i++){
  selectWindow(""+i);
  close();
}
selectWindow("Result of 0");
close();
selectWindow("Result of Result of 0");
rename(a);
// Colour Thresholding-------------
run("Analyze Particles...", "size=3000-Infinity include summarize");
close();
print("total_area");
selectWindow("Summary");
lines = split(getInfo(), "\n");
headings = split(lines[0], "\t");
values = split(lines[lengthOf(lines)-1], "\t");
 for (i=0; i<headings.length; i++)
     print(headings[i]+": "+values[i]);
} else
            print("Error opening "+path);
//////////////////////////////////////////////yellow area
showProgress(k, files.length);
        IJ.redirectErrorMessages();
```

```
        open(path);
        if (nImages>=1) {
//remove background
setBackgroundColor(0, 0, 0);
run("Clear Outside");
// Color Thresholder 1.51j
// Autogenerated macro, single images only!
min=newArray(3);
max=newArray(3);
filter=newArray(3);
a=getTitle();
run("HSB Stack");
run("Convert Stack to Images");
selectWindow("Hue");
rename("0");
selectWindow("Saturation");
rename("1");
selectWindow("Brightness");
rename("2");
min[0]=0;
max[0]=43;
filter[0]="pass";
min[1]=0;
max[1]=255;
filter[1]="pass";
min[2]=164;
max[2]=255;
filter[2]="pass";
filter[0]="pass";
filter[1]="pass";
filter[2]="pass";
for (i=0;i<3;i++){
  selectWindow(""+i);
  setThreshold(min[i], max[i]);
  run("Convert to Mask");
  if (filter[i]=="stop")  run("Invert");
}
imageCalculator("AND create", "0","1");
imageCalculator("AND create", "Result of 0","2");
for (i=0;i<3;i++){
  selectWindow(""+i);
  close();
}
selectWindow("Result of 0");
close();
selectWindow("Result of Result of 0");
```

```
rename(a);
// Colour Thresholding------------
run("Analyze Particles...", "size=10-Infinity include summarize");
close();
print("yellow");
selectWindow("Summary");
lines = split(getInfo(), "\n");
headings = split(lines[0], "\t");
values = split(lines[lengthOf(lines)-1], "\t");
 for (i=0; i<headings.length; i++)
      print(headings[i]+": "+values[i]);
} else
            print("Error opening "+path);
////////////////////////////////////////////black area
showProgress(k, files.length);
        IJ.redirectErrorMessages();
        open(path);
        if (nImages>=1) {
//remove background
setBackgroundColor(0, 0, 0);
run("Clear Outside");
// Color Thresholder 1.51j
// Autogenerated macro, single images only!
min=newArray(3);
max=newArray(3);
filter=newArray(3);
a=getTitle();
run("HSB Stack");
run("Convert Stack to Images");
selectWindow("Hue");
rename("0");
selectWindow("Saturation");
rename("1");
selectWindow("Brightness");
rename("2");
min[0]=0;
max[0]=33;
filter[0]="pass";
min[1]=0;
max[1]=255;
filter[1]="pass";
min[2]=1;
max[2]=155;
filter[2]="pass";
filter[0]="pass";
filter[1]="pass";
```

```
filter[2]="pass";
for (i=0;i<3;i++){
  selectWindow(""+i);
  setThreshold(min[i], max[i]);
  run("Convert to Mask");
  if (filter[i]=="stop")  run("Invert");
}
imageCalculator("AND create", "0","1");
imageCalculator("AND create", "Result of 0","2");
for (i=0;i<3;i++){
  selectWindow(""+i);
  close();
}
selectWindow("Result of 0");
close();
selectWindow("Result of Result of 0");
rename(a);
// Colour Thresholding-------------
run("Analyze Particles...", "size=20-Infinity include summarize");
close();
print("black");
selectWindow("Summary");
lines = split(getInfo(), "\n");
headings = split(lines[0], "\t");
values = split(lines[lengthOf(lines)-1], "\t");
 for (i=0; i<headings.length; i++)
     print(headings[i]+": "+values[i]);
} else
             print("Error opening "+path);
//////////////////////////////////////////////black_dot
showProgress(k, files.length);
        IJ.redirectErrorMessages();
        open(path);
        if (nImages>=1) {

//remove background
setBackgroundColor(0, 0, 0);
run("Clear Outside");

// Color Thresholder 1.51j
// Autogenerated macro, single images only!
min=newArray(3);
max=newArray(3);
filter=newArray(3);
a=getTitle();
run("HSB Stack");
```

```
run("Convert Stack to Images");
selectWindow("Hue");
rename("0");
selectWindow("Saturation");
rename("1");
selectWindow("Brightness");
rename("2");
min[0]=0;
max[0]=33;
filter[0]="pass";
min[1]=0;
max[1]=255;
filter[1]="pass";
min[2]=1;
max[2]=155;
filter[2]="pass";
filter[0]="pass";
filter[1]="pass";
filter[2]="pass";
for (i=0;i<3;i++){
  selectWindow(""+i);
  setThreshold(min[i], max[i]);
  run("Convert to Mask");
  if (filter[i]=="stop")  run("Invert");
}
imageCalculator("AND create", "0","1");
imageCalculator("AND create", "Result of 0","2");
for (i=0;i<3;i++){
  selectWindow(""+i);
  close();
}
selectWindow("Result of 0");
close();
selectWindow("Result of Result of 0");
rename(a);
// Colour Thresholding-------------

run("Analyze Particles...", "size=20-1000 circularity=0.30-1.00 show=Nothing include
summarize");
close();
print("black_dot");
selectWindow("Summary");
lines = split(getInfo(), "\n");
headings = split(lines[0], "\t");
values = split(lines[lengthOf(lines)-1], "\t");
 for (i=0; i<headings.length; i++)
```

```
      print(headings[i]+": "+values[i]);
} else
            print("Error opening "+path);
//////////////////////////////////////////////green area
showProgress(k, files.length);
        IJ.redirectErrorMessages();
        open(path);
        if (nImages>=1) {
//remove background
setBackgroundColor(0, 0, 0);
run("Clear Outside");
// Color Thresholder 1.51j
// Autogenerated macro, single images only!
min=newArray(3);
max=newArray(3);
filter=newArray(3);
a=getTitle();
run("HSB Stack");
run("Convert Stack to Images");
selectWindow("Hue");
rename("0");
selectWindow("Saturation");
rename("1");
selectWindow("Brightness");
rename("2");
min[0]=37;
max[0]=135;
filter[0]="pass";
min[1]=42;
max[1]=255;
filter[1]="pass";
min[2]=1;
max[2]=255;
filter[2]="pass";
filter[0]="pass";
filter[1]="pass";
filter[2]="pass";
for (i=0;i<3;i++){
  selectWindow(""+i);
  setThreshold(min[i], max[i]);
  run("Convert to Mask");
  if (filter[i]=="stop")  run("Invert");
}
imageCalculator("AND create", "0","1");
imageCalculator("AND create", "Result of 0","2");
for (i=0;i<3;i++){
```

```
  selectWindow(""+i);
  close();
}
selectWindow("Result of 0");
close();
selectWindow("Result of Result of 0");
rename(a);
// Colour Thresholding------------
run("Analyze Particles...", "size=0-Infinity circularity=0.30-1.00 show=Nothing include
summarize");
close();
print("green");
selectWindow("Summary");
lines = split(getInfo(), "\n");
headings = split(lines[0], "\t");
values = split(lines[lengthOf(lines)-1], "\t");
 for (i=0; i<headings.length; i++)
     print(headings[i]+": "+values[i]);
} else
           print("Error opening "+path);
}
```

**Heritability Equations**

Severity from the detached leaf (DTVI and DTIA)

$$H_{DT}^2 = \frac{\sigma_g^2}{\sigma_g^2 + \frac{\sigma_e^2}{rep * \exp} + \frac{\sigma_{ge}^2}{exp} + \frac{\sigma_{gp}^2}{plate}}$$

Severity from the leaf disk assay (DSVI and DSIA)

$$H_{DS}^2 = \frac{\sigma_g^2}{\sigma_g^2 + \frac{\sigma_e^2}{rep * plate} + \frac{\sigma_{ge}^2}{plate}}$$

Severity in field from two locations

$$H_{TL}^2 = \frac{\sigma_g^2}{\sigma_g^2 + \sigma_e^2}$$

Severity in field from NY

$$H_{NY}^2 = \frac{\sigma_g^2}{\sigma_g^2 + \sigma_e^2}$$

Severity in field from PA

$$H_{PA}^2 = \frac{\sigma_g^2}{\sigma_g^2 + \sigma_e^2}$$

$\sigma_g^2 = variance\ due\ to\ genetic$

$\sigma_e^2 = variance\ due\ to\ residual\ error$

$\sigma_{ge}^2 = variance\ due\ to\ interaction\ between\ genetics\ and\ environment\ like\ experiment$

$\sigma_{gp}^2 = variance\ due\ to\ interaction\ between\ genetics\ and\ plate\ in\ leaf\ disk\ assay$

$rep = number\ of\ replicates$

$\exp = number\ of\ experiment\ set$

**QQ-plots**

In the full set of 478 genotypes, single traits from artificial inoculation via either detached leaf or leaf disk assay with vision or image analysis (DTVI, DTIA, DSVI, and DSIA) provided significant GWAS results based on inflated QQ-plot (Figure S5-6). Although DSVI and DSIA showed the best QQ-plot (Figure S5c-d), the convex head and inflated tail of -log(P-value) suggested confounding of the association analysis. The single-trait GWAS of field evaluations (MTL, MNY, MPA, BNY, BPA, mean TL, mean of NY, and mean of PA) did not yield any significant peaks from association analysis (Figure S6). The combination GWAS of two traits among DTVI, DTIA, DSVI, and DSIA did not provide more significant peaks despite than single traits (Figure S7a-c). The combination of three traits of DTVI, DTIA and DSIA providing the most significant peaks and theoretically expected QQ-plot (Figure S7d).

In a tetraploid group, the same three-trait combination of DTVI-DTIA-DSIA did not provide the better GWAS than did DSVI-MNY (Figure S8d). The GWAS from DSVI and MNY separately seemed to provide significant peaks but QQ-plot was not distributed as in theory. Similarly, in octaploid, DTVI-DTIA-DSIA combination did provide the same GWAS profile as DTVI-DSIA combination (Figure S9). Although DSIA seemed to provide significant peaks, QQ-plot was not distributed as in theory.

None of the image analysis traits and combination with BLUPs from NY and PA provided a significant peak from GWAS in lowland (Figure S10). Also, the upland combination of DTVI-DTIA-DSIA did not yield significant peaks from GWAS (Figure S11).
